# MYPT1 O-GlcNAc modification controls the sensitivity of fibroblasts to sphingosine-1-phosphate mediated cellular contraction

**DOI:** 10.1101/796672

**Authors:** Nichole J. Pedowitz, Anna R. Batt, Narek Darabedian, Matthew R. Pratt

## Abstract

Many intracellular proteins can be modified by N-acetylglucosamine, a posttranslational modification known as O-GlcNAc. Because this modification is found on serine and threonine side-chains, O-GlcNAc has the potential to dynamically regulate cellular signaling pathways through interplay with phosphorylation. Here, we discover and characterize one such example. First, we find that O-GlcNAc levels control the sensitivity of fibroblasts to actin contraction induced by the signaling lipid sphingosine-1-phosphate (S1P). In follow-up mechanistic investigations, we show that this O-GlcNAc dependence lies in the signaling pathway through the S1PR2 receptor and subsequent activation of the Rho and Rho kinase. This pathway typically culminates in the phosphorylation of myosin light chain (MLC), resulting in myosin activation and cellular contraction. We discovered that O-GlcNAc modification of the phosphatase subunit MYPT1 inhibits this pathway by blocking MYPT1 phosphorylation, maintaining its activity and causing the dephosphorylation of MLC. Therefore, MYTP1 O-GlcNAc levels function to regulate the sensitivity of cells to S1P-mediated cellular contraction. Finally, we demonstrate that O-GlcNAc levels alter the sensitivity of primary human dermal fibroblasts in a collagen matrix model of wound healing. Our findings have important implications for the role of O-GlcNAc in fibroblast motility and differentiation, particularly in diabetic wound healing, where increased levels of the modification may inhibit S1P-mediated healing phenotypes in fibroblasts.

## INTRODUCTION

O-GlcNAc is a form of protein glycosylation that is uniquely suited to regulate cellular signaling pathways (Figure 1a). This intracellular posttranslational modification involves the transfer of the monosaccharide *N*-acetylglucosamine to serine and threonine side chains of proteins throughout the cytosol, nucleus, and mitochondria of all animal cells^1–3^. Unlike most forms of cell surface glycosylation, this core GlcNAc moiety is not further elaborated by additional carbohydrates and is dynamically regulated through the action of two enzymes. O-GlcNAc transferase (OGT) uses the donor sugar UDP-GlcNAc to install O-GlcNAc onto substrates, while O-GlcNAcase (OGA) removes it^4–6^, setting up a scenario potentially similar to the cycling of protein phosphorylation. Additionally, the amounts of O-GlcNAc have been shown to change upon alterations in cellular environment through the sensing of different biological inputs. For example, one of the main roles of O-GlcNAc is to act as a nutrient sensor to control various cellular pathways^7^. In particular, flux of glucose into the cell, which is transformed by the hexosamine biosynthetic pathway (HBP) into increased amounts of UDP-GlcNAc^8^, results in a dynamic increase in overall O-GlcNAc levels. Interestingly, the substrate selection and activity of OGT are highly dependent on the concentration of UDP-GlcNAc^9^. This enzymatic feature of OGT means that O-GlcNAcylation of proteins is a “read-out” of extracellular glucose levels. Therefore, it is not surprising that hyperglycemia in diabetes results in elevated O-GlcNAc in comparison to healthy individuals. Evidence from human genetics also supports an important role for increased O-GlcNAc in the development of diabetes. Specifically, polymorphisms in the gene encoding OGA result in elevated O-GlcNAc an increased risk for type II diabetes and a lower age of onset^10^. Taken together, these general observations indicate that O-GlcNAc should modulate various proteins and pathways and that misregulation of these events contributes to human disease. An attractive hypothesis in the O-GlcNAc field has been a direct relationship between the O-GlcNAc modification and phosphorylation of proteins (Figure 1b) ^11^. For example, O-GlcNAc modification of the kinase Akt at Thr305 and Thr312 prevents phosphorylation at nearby Thr308 and impairs Akt signaling^12^. Similarly, O-GlcNAc modification of casein kinase II (CK2) at Ser347 antagonizes phosphorylation at nearby Thr344, thereby reducing the stability and activity of this kinase^13^. In contrast, activating phosphorylation of the transcription factor cyclic AMP-response element binding protein (CREB) at Ser133 is required for subsequent O-GlcNAc at Ser40, which can then repress its transcriptional activity with key consequences for memory formation^14^. However, despite these and some other examples, and the theoretical attractiveness of O-GlcNAc as a regulator of cell signaling, the cases where we understand how this modification controls specific cascades remains limited.

**Figure 1.**
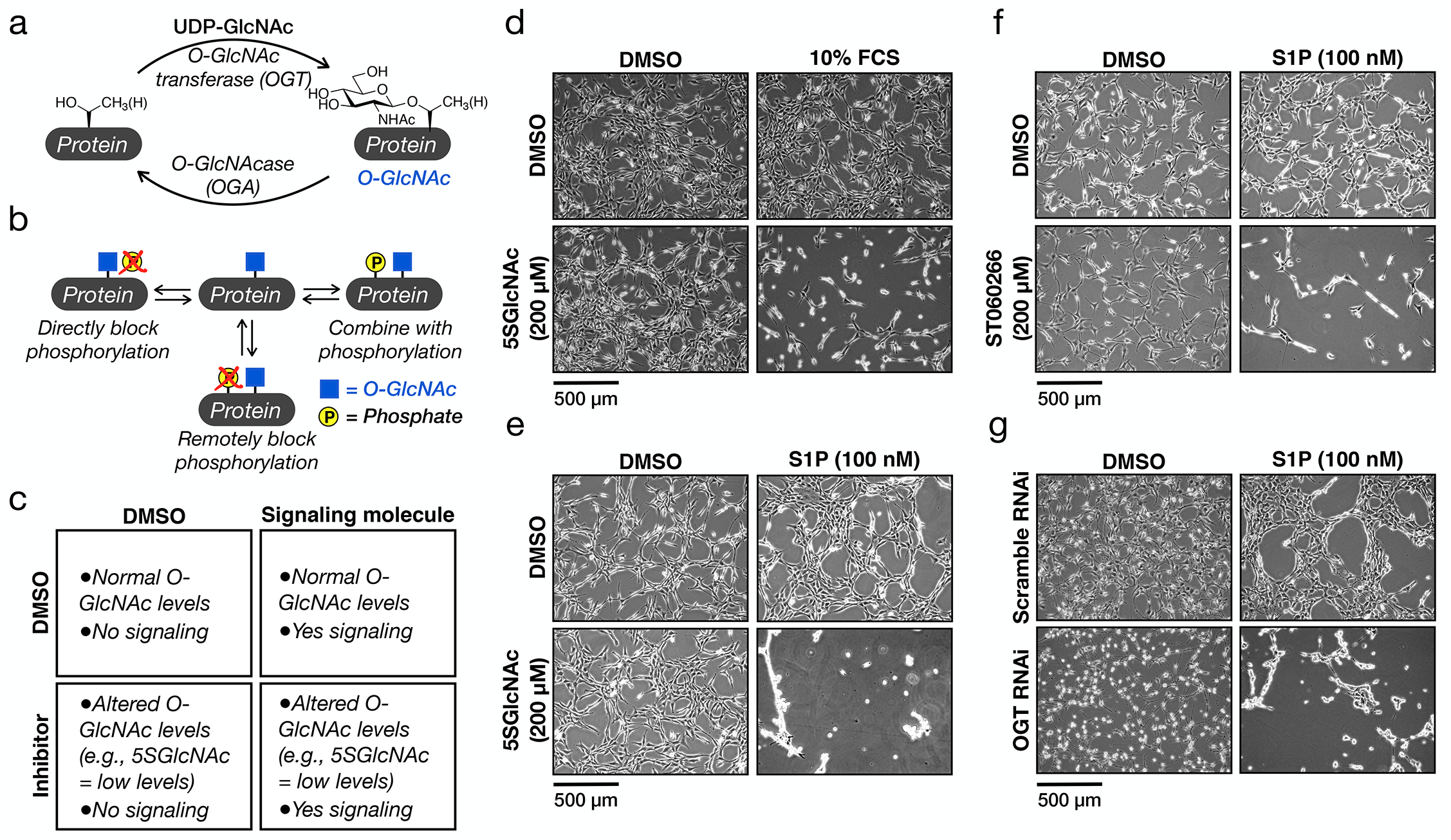
O-GlcNAc controls the sensitivity of fibroblasts to sphingosine-1-phosphate (S1P) mediated cell contraction. a) O-GlcNAcylation is the dynamic addition to N-acetylglucosamine to serine and threonine residues of intracellular proteins. b) O-GlcNAcylation has its own functions but can also antagonize phosphorylation, both directly and remotely, as well as combine with phosphorylation to illicit biological outcomes. c) Schematic layout of our general experimental conditions. d) NIH3T3 cells were treated with either DMSO or the OGT inhibitor 5SGlcNAc (200 μM) for 16 h before addition of either more DMSO or 10% FCS for 30 min. The contraction phenotype was then visualized using bright-field microscopy. e) NIH3T3 cells were treated with either DMSO or 5SGlcNAc (200 μM) for 16 h before addition of either more DMSO or S1P (100 nM) for 30 min. The contraction phenotype was then visualized using bright-field microscopy. f) NIH3T3 cells were treated with either DMSO or the OGT inhibitor ST060266 (200 μM) for 16 h before addition of either more DMSO or S1P (100 nM) for 30 min. The contraction phenotype was then visualized using bright-field microscopy. g) NIH3T3 cells were transfected with either scramble or OGT-targeted RNAi for 48 h before addition of either more DMSO or S1P (100 nM) for 30 min. The contraction phenotype was then visualized using bright-field microscopy.

One reason for our narrow understanding of O-GlcNAc has been an absence of applicable biological tools. Genetic ablation of either OGT or OGA is lethal during development in most model organisms and loss of OGT results in mammalian cell death in culture^15,16^. Additionally, while antibodies against O-GlcNAc are available they do not detect all modifications nor can be readily used to determine absolute O-GlcNAc stoichiometry. Fortunately, a range of chemical tools have been developed over the past decade that overcome these limitations, including small molecule inhibitors of both OGT^17^ and OGA^18^ that enable the dynamic control of overall O-GlcNAc levels and chemoenzymatic labeling techniques^19,20^ for the identification and characterization of protein substrates. Here, we apply these chemical tools to investigate our serendipitous discovery linking the signaling lipid sphingosine-1-phosphate (S1P) to actin cytoskeletal remodeling through the O-GlcNAc modification status of the phosphatase regulatory subunit MYPT1.

## RESULTS

### O-GlcNAc controls fibroblast contraction in response to the serum lipid sphingosine-1-phosphate

During the course of our previous investigation into the roles of O-GlcNAc in apoptosis, we encountered a dramatic phenotypic change in NIH3T3 fibroblasts upon reduction of overall O-GlcNAc levels followed by treatment with fetal calf serum (FCS). As schematized in Figure 1c, these fibroblasts were treated with the OGT inhibitor 5SGlcNAc (200 μM) for 16 h to give a downregulation of global O-GlcNAc (Supplementary Figure 1a). Upon subsequent addition of 10% FCS, the cells underwent a rapid (10 - 30 min) morphological change, resulting in their contraction on the culture plate (Figure 1d and Supplementary Video 1). Intrigued by this result, we then screened multiple protein factors, cytokines, and lipids commonly found in serum. Strikingly, we only detected this phenotype upon S1P treatment (100 nM), and to a lesser extent lysophosphatidic acid (LPA) treatment (20 μM), but not with any of the other serum components (Figure 1e, Supplementary Figure 1e and Supplementary Video 2), indicating that cellular signaling initiated by these lipids is responsible for our observation. Given the difference in potency between these two lipids, we chose to explore S1P in greater detail. In order to confirm that this phenotype is indeed due to a reduction in O-GlcNAc and not a potential off-target effect of 5SGlcNAc, we next treated fibroblasts with either an orthogonal OGT inhibitor (ST060266, 200 μM) or RNAi directed again OGT (Supplementary Figure 1b & c). In both cases, we found a similar morphological change upon addition of S1P (Figures 1f & g), confirming the O-GlcNAc dependence of the phenotype. To test whether this phenotype was apoptosis, we next used Western blotting to visualize any cleavage of caspase-3 during 5SGlcNAc and/or S1P treatment and did not observe any decrease in the full-length protein or appearance of the active protease (Supplementary Figure 2). Next, we examined whether the cells would return to a relaxed state. Specifically, we treated NIH3T3 cells with with DMSO or 5SGlcNAc (200 μM) followed by S1P (100 nM) for an extended, 180 min, time course (Supplementary Figure 3). Again, we found no morphological change in the cells treated only with S1P. However, consistent with our previous observations, the cells with decreased O-GlcNAc levels quickly contracted but returned to a more relaxed and attached state over the course of the assay. Together, these data suggest that O-GlcNAc controls the sensitivity of fibroblasts to a dynamic, S1P-mediated signaling pathway.

Notably, published studies previously demonstrated that μM concentrations of S1P cause fibroblast contraction in collagen matrices^21,22^, strongly suggesting that we are increasing the sensitivity of this same pathway by lowering O-GlcNAc levels. To test this possibility, we again treated NIH3T3 cells with either DMSO vehicle or 5SGlcNAc (200 μM) for 16 h followed by a range of S1P concentrations from 0.05 to 5 μM. Consistent with published results^22^, fibroblasts with “normal” O-GlcNAc levels underwent contraction at the higher concentrations of S1P, but cells treated with the OGT inhibitor displayed increased sensitivity to lower amounts of S1P (Figure 2a). We then quantified the amounts of cellular contraction by measuring the surface area occupied by the cells and found the difference in contraction to be statistically significant (Figure 2b). Importantly, this increased sensitivity was observed in multiple biological replicates (Supplementary Figure 4a). Given that NIH3T3 cells with baseline O-GlcNAc will respond to the higher concentrations of S1P, we next asked whether raising the modification levels would decrease their sensitivity to the signaling lipid. Accordingly, we treated these fibroblasts with either DMSO vehicle or the OGA inhibitor Thiamet-G (10 μM), resulting in increased overall O-GlcNAc (Supplementary Figure 1a). Gratifyingly, we found the Thiamet-G treated cells to be significantly more resistant to S1P concentrations compared to the control (Figures 2c & d), and again this phenotypic difference was recapitulated in multiple biological replicates (Supplementary Figures 4b).

**Figure 2.**
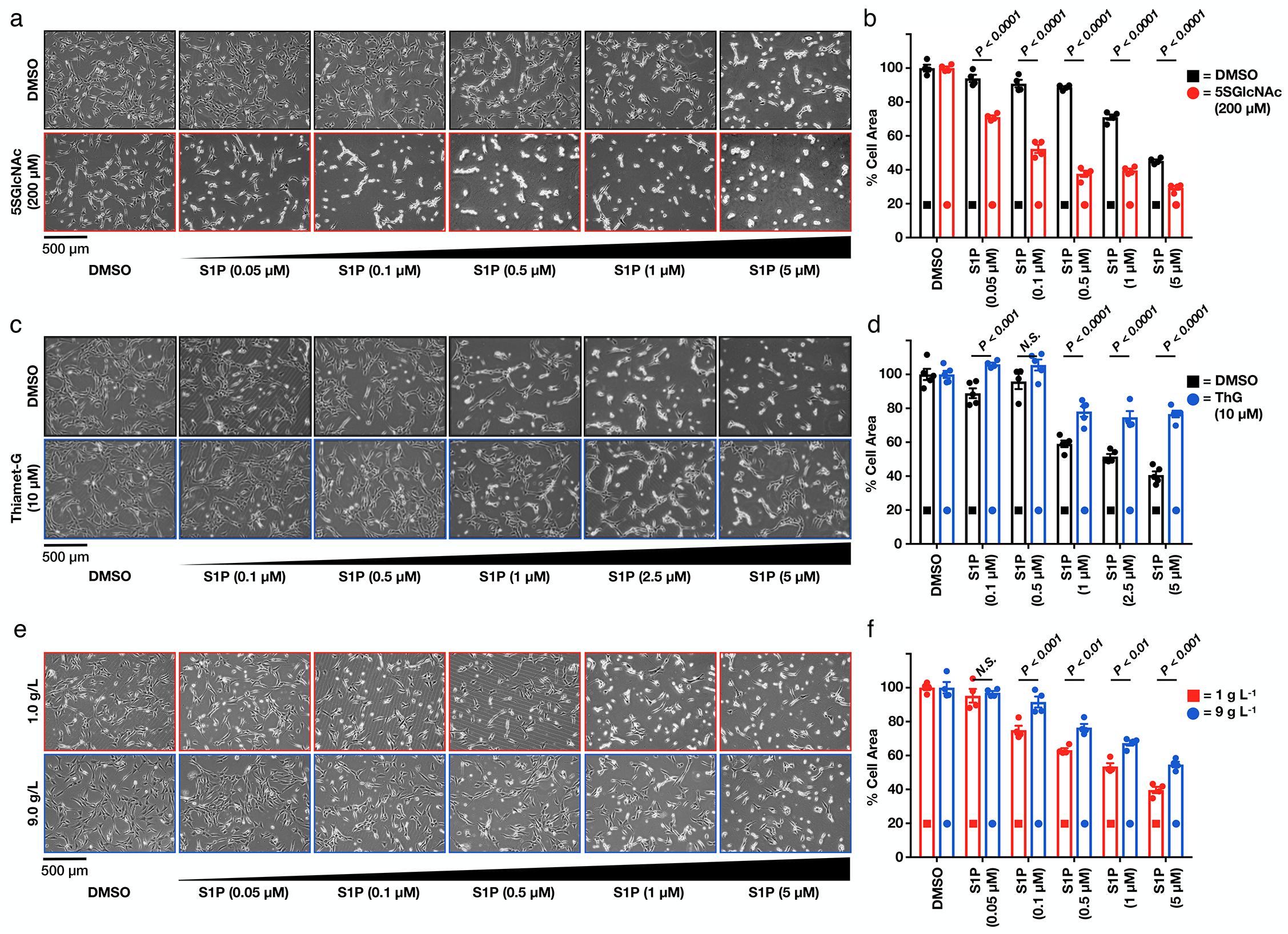
O-GlcNAc controls the sensitivity of cells to S1P-mediated contraction. a) Lowering O-GlcNAc levels increases the sensitivity of NIH3T3 cells to S1P induced cell contraction. NIH3T3 cells were pre-treated with either DMSO or 5SGlcNAc (200 μM) for 16 h before addition of the indicated concentrations of S1P for 30 min. The contraction phenotype was then visualized using bright-field microscopy. b) Quantitation of the data in (a). Results are the mean ± SEM of the relative culture plate area taken-up by cells in four randomly selected frames. Statistical significance was determined using a 2-way ANOVA test followed by Sidak’s multiple comparisons test. c) Raising O-GlcNAc levels decreases the sensitivity of NIH3T3 cells to S1P induced cell contraction. NIH3T3 cells that had been treated with either DMSO or the OGA inhibitor Thiamet-G (10 μM) for 20 h before addition of the indicated concentrations of S1P for 30 min. The contraction phenotype was then visualized using bright-field microscopy. d) Quantitation of the data in (c). Results are the mean ± SEM of the relative culture plate area taken-up by cells in four randomly selected frames. Statistical significance was determined using a 2-way ANOVA test followed by Sidak’s multiple comparisons test. e) Glucose concentration controls the sensitivity of NIH3T3 cells to S1P induced cell contraction. NIH3T3 cells were cultured in the indicated amounts of glucose for 48 h before addition of the indicated concentrations of S1P for 30 min. The contraction phenotype was then visualized using bright-field microscopy. f) Quantitation of the data in (e). Results are the mean ± SEM of the relative culture plate area taken-up by cells in four randomly selected frames. Statistical significance was determined using a 2-way ANOVA test followed by Sidak’s multiple comparisons test.

As mentioned in the introduction, O-GlcNAc modifications rise due to higher levels of circulating glucose levels, with important consequences in diabetes. This observation and others raised the possibility that the modification could act as a nutrient sensor to control S1P-dependent signaling. To test this possibility, we cultured NIH3T3 cells in media containing either 1 or 9 g L^−1^ glucose for 48 h, resulting in the expected differences in O-GlcNAc levels (Supplementary Figure 1d). As predicted, we found that the cells cultured in low glucose were indeed more sensitive to S1P-mediated contraction compared to those grown in high glucose (Figures 2e & f and Supplementary Figures 4c). Together these results indicate that O-GlcNAcylation acts as a molecular buffer to control S1P signaling that culminates in fibroblast contraction.

### The O-GlcNAc-dependent, S1P-mediated contraction phenotype results from signaling through one of five S1P G-protein coupled receptors (S1PR2)

S1P is an endogenous signaling lipid^23^ produced intracellularly from the phosphorylation of ceramide by sphingosine kinases 1 and 2 (SphK1, SphK2). It can then be transported outside of the cell and signal through a family of five G-protein coupled receptors (GPCRs) (S1PR1 through S1PR5, Supplementary Figure 5). Subsequent coupling of these GPCRs to different classes of G-proteins results in the activation of several associated downstream signaling pathways^24–26^. S1P plays roles in many biological events, including cell survival, proliferation, motility, and adherence. Therefore, it is not surprising that it has been shown to contribute to diverse areas of human health and disease, such as cardiovascular development and control of blood pressure, regulation of immune cell migration, and wound healing^27–30^.

As a next step towards discovering the molecular mechanism by which O-GlcNAc regulates S1P signaling, we first used RT-PCR to determine which of the S1PRs were actively transcribed in our fibroblasts. We found that all five receptors had the potential to be expressed in this cell line (Supplementary Figure 6a). Given that this approach was unable to narrow down the potential S1PR responsible, we then turned to small molecule pharmacology. Specifically, we first treated NIH3T3 cells with either DMSO or 5SGlcNAc (200 μM) to modulate O-GlcNAc levels as above. We then added individual selective antagonists of S1PR1 through S1PR4 (1 μM) for 10 min immediately followed by S1P (100 nM). Visualization of the cellular contraction phenotype demonstrated that only blockage of S1PR2 signaling could prevent the S1P-mediated morphological change (Supplementary Figures 6b & c), indicating that the other three receptors (S1PR1, R3, and R4) are not involved. Unfortunately, S1PR5 does not yet have a selective antagonist, and therefore we could not rule it out with these data. However, when we treated NIH3T3 cells with a selective agonist of this receptor, we observed no phenotypic change regardless of O-GlcNAc levels (Supplementary Figures 6d & e). Next, to test whether S1PR2 agonism alone is sufficient to activate the pathway, we exposed DMSO- or 5SGlcNAc-treated cells to different concentrations of a selective S1PR2 agonist. Notably, we found that low concentrations of this agonist (1 μM) resulted in cell contraction under low O-GlcNAcylation levels while a higher concentration (5 μM) induced the phenotype universally (Supplementary Figures 6f & g), similar to what we observed for different concentrations of S1P. Finally, we knocked down S1PR2 using RNAi and found that S1P (100 nM) did not induce cell contraction under either normal or reduced O-GlcNAc levels (Supplementary Figures 6h & i). Together these results demonstrate that O-GlcNAcylation controls a signaling pathway that is likely downstream of S1PR2.

### O-GlcNAc regulates the activation of myosin light chain (MLC) and its associated phosphatase MYPT1 in response to S1P

Previous work has shown that stressed collagen-matrix contraction by fibroblasts is dependent on Rho kinase^21^. Therefore, we next decided to focus on S1P signaling through Rho GTPase, where S1PR agonism results in activation of Rho and Rho kinase (ROCK1/2) ^31^, ultimately giving actin contraction (Figure 3a and Supplementary Figure 5) ^25,32–34^. More specifically, activated ROCK1/2 will phosphorylate myosin light chain (MLC) at two serines (Thr18 & Ser19) resulting in the formation of an active myosin complex and actin filament contraction^34–36^. To avoid unwanted activation and terminate this pathway, cells are equipped with molecular brakes in the form of the phosphatase regulatory subunit MYPT1 that mediates the dephosphorylation MLC. Previous work has shown that ROCK1/2 can also phosphorylate MYTP1 at Thr696 and Thr853^37–40^. This results in the deactivation of MYPT1^41^, effectively cutting the breaks on the pathway and ensuring myosin activation.

**Figure 3.**
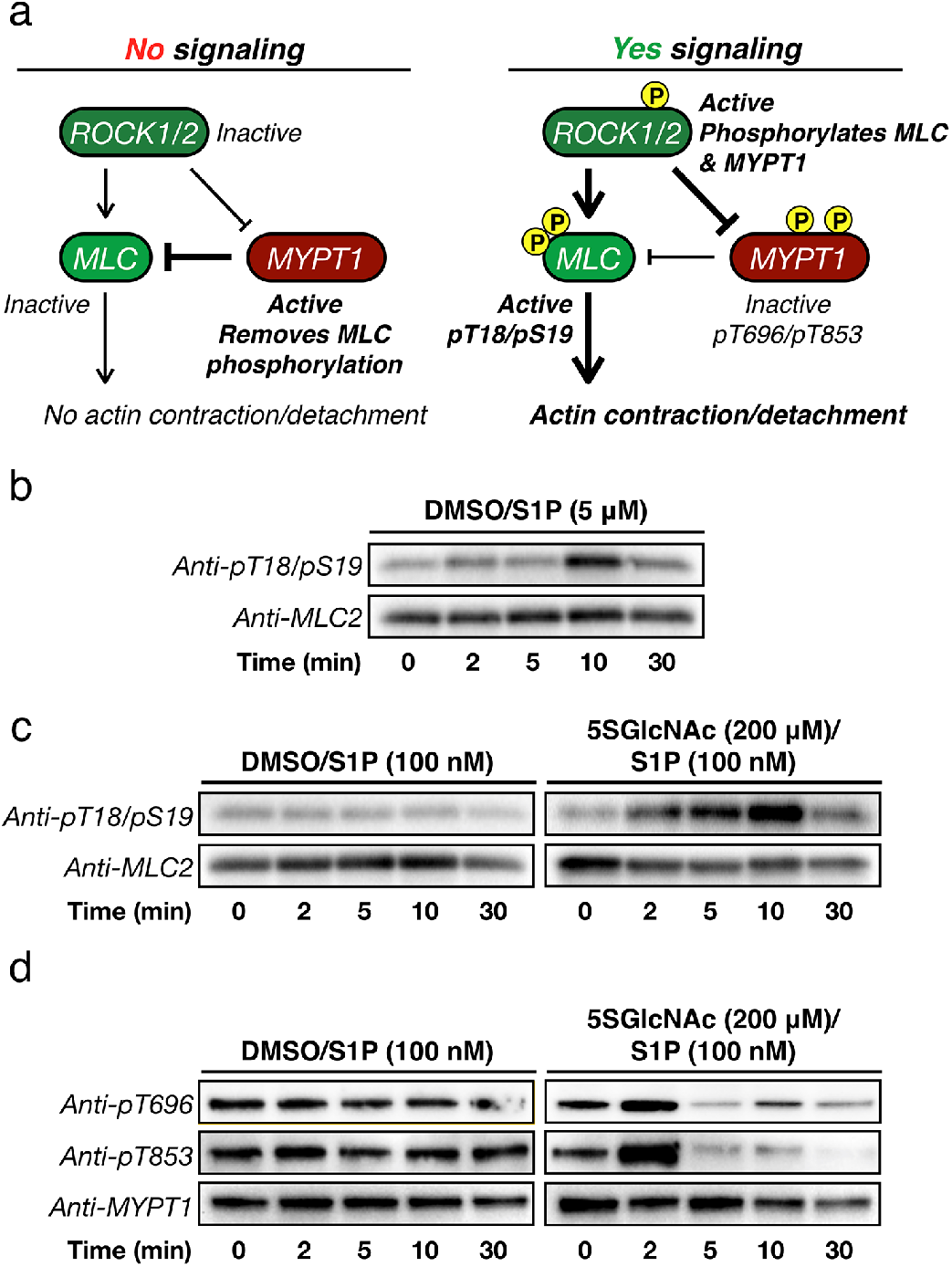
O-GlcNAc levels control S1P-mediated phosphorylation of MLC and MYPT1. a) Schematic of the ROCK-signaling pathway that can result in cellular contraction. Under no or low signaling conditions ROCK does not phosphorylate MLC or MYPT1, and MYPT1 remains active to dephosphorylate MLC. Upon induction of signaling, ROCK phosphorylates both MLC to activate contraction and MYPT1 to inactivate dephosphorylation, effectively pushing on the gas and cutting the brake. b) MLC is phosphorylated under high S1P signaling. NIH3T3 cells were treated with S1P (5 μM) for the indicated lengths of time. MLC phosphorylation, and thus activation, was visualized using Western blotting. c) MLC is phosphorylated under low S1P signaling when O-GlcNAc levels are reduced. NIH3T3 cells were treated with either DMSO or 5SGlcNAc (200 μM) for 16 h followed by S1P (100 nM) for the indicated lengths of time. MLC phosphorylation, and thus its activation, was visualized using Western blotting. d) MYPT1 is phosphorylated under low S1P signaling when O-GlcNAc levels are reduced. NIH3T3 cells were treated with either DMSO or 5SGlcNAc (200 μM) for 16 h followed by S1P (100 nM) for the indicated lengths of time. MYPT1 phosphorylation, and thus its deactivation, was visualized using Western blotting.

To test whether ROCK1/2 activity was indeed required for our observed contraction phenotype, we again treated NIH3T3 cells with either DMSO or 5SGlcNAc (200 μM) for 16 h, followed by the ROCK1/2 inhibitor Y27632 (10 μM) for an additional hour before addition of S1P (0.05 - 5 μM). Notably, we observed no contraction of any cells at all concentrations of S1P (Supplementary Figure 7). With this result confirming the importance of ROCK1/2, we moved on to explore the phosphorylation status of its downstream targets MLC^42^ and MYPT1^43,44^. In order to confirm these previously published results, we first treated NIH3T3 cells with normal O-GlcNAcylation levels to a high concentration of S1P (5 μM) that results in their contraction. After different lengths of time, we collected the cells and subjected the corresponding lysates to analysis by Western blotting (Figure 3b). As expected, we observed increasing phosphorylation of MLC at Ser18/Thr19 that corresponded well with the overall timing of cell contraction. Next, we performed a similar analysis of NIH3T3 cells that had been pre-treated with either DMSO or 5SGlcNAc (200 μM) for 16 h before addition of a low concentration of S1P (100 nM) (Figure 3c & d). In cells treated with DMSO and thus having normal O-GlcNAcylation levels, we found no detectable increase in MLC or MYTP1 phosphorylation. However, in cells with depleted O-GlcNAc (i.e., 5SGlcNAc treated), we detected a rapid increase in phosphorylation of both MLC and MYPT1. Importantly, the oscillation of phosphorylation on MYTP1 that we see is consistent with previous reports and relates to the function of MYTP1 as its own phosphatase and therefore a biological timer of this signaling pathway in response to compressive force^45^.

### MYPT1 is a heavily and dynamically O-GlcNAc modified protein

We next set out to identify which protein in the Rho/ROCK signaling pathway is potentially regulated by O-GlcNAc. We chose to focus on MLC and MYPT1 first, as they were both some of the earliest O-GlcNAc modified proteins identified^46,47^. In order to ascertain the O-GlcNAcylation status of these proteins, we took advantage the chemoenzymatic labeling protocol originally developed by the Hsieh-Wilson and Qasba labs (Figure 4a) ^19,48^. Briefly, we first used this method to install a cleavable biotin-tag onto all endogenous O-GlcNAc modifications in NIH3T3 cell lysates. After incubation with streptavidin beads and extensive washing, the O-GlcNAcylated proteins were eluted and analyzed by Western blotting. In accordance with previous reports, we identified both MLC and MYPT1 to be O-GlcNAc modified (Figure 4b). The known O-GlcNAc modified protein nucleoporin 62 (Nup62) served as a positive control with β-actin as a negative control in this experiment. Notably, while both MLC and MYPT1 are indeed O-GlcNAc modified, when we normalized the amount of input to our O-GlcNAc pulldown, we found that MYPT1 was modified at dramatically higher levels compared to MLC and similar to the constitutively O-GlcNAc-modified Nup62 (Figure 4c). Therefore, we chose to move forward with MYPT1 as the most likely candidate to be regulated in the pathway.

**Figure 4.**
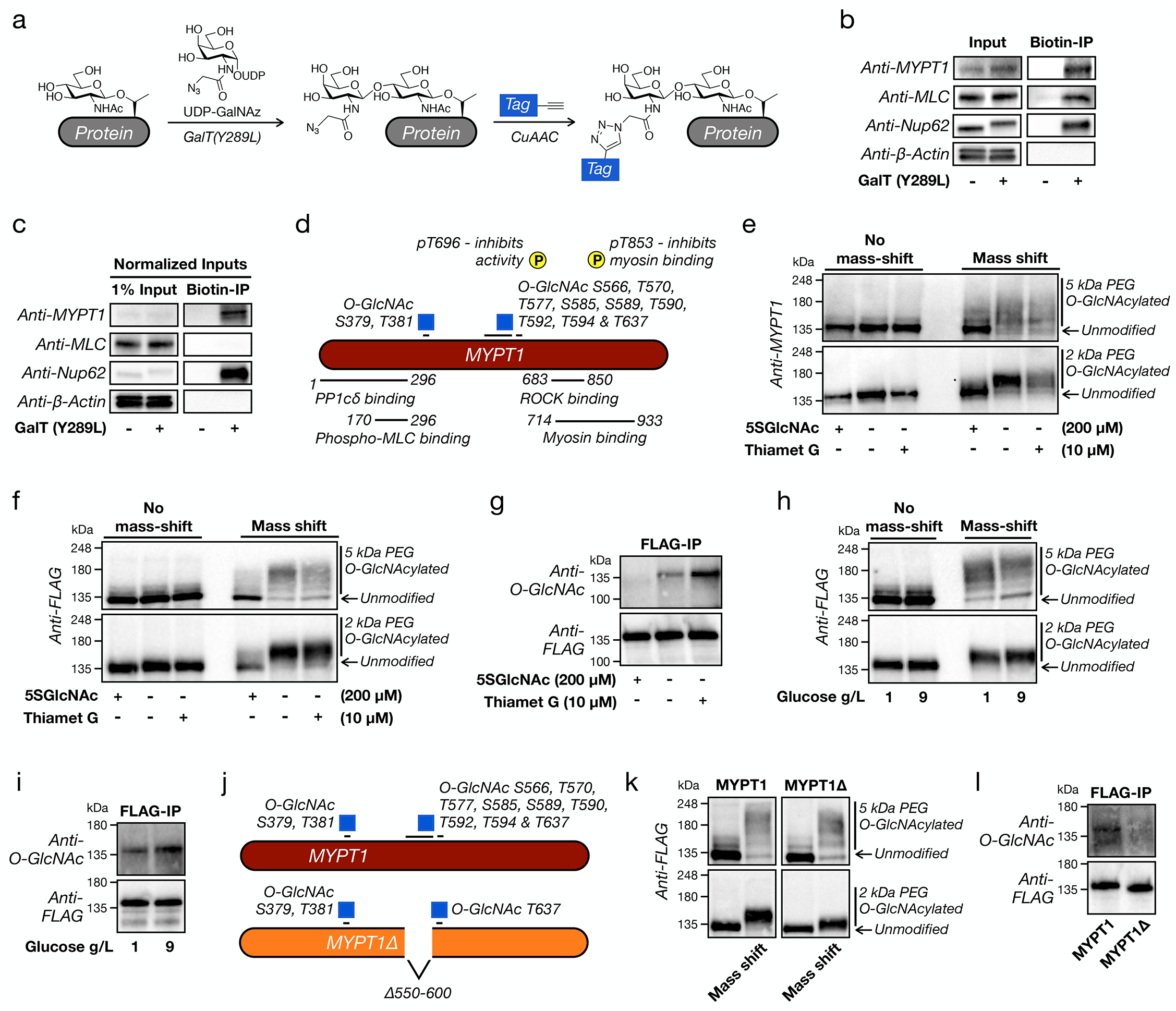
MYPT1 is heavily and dynamically O-GlcNAc modified near its ROCK-binding domain. a) The chemoenzymatic method for detecting O-GlcNAc modifications. Endogenous O-GlcNAc moieties are enzymatically modified with an azide-containing sugar that can then be used for the subsequent installation of tags using bioorthogonal chemistry. b) MLC and MYPT1 are both O-GlcNAc modified. O-GlcNAc modified proteins from NIH3T3 cells were enriched using chemoenzymatic labelling and analyzed by Western blotting. Nucleoporin 62 (Nup62) is a known, heavily O-GlcNAc modified protein, and β-actin serves as a negative control. c) MYPT1 is highly O-GlcNAc modified, while MLC is not. the samples in (b) were normalized for inputs and analyzed by Western blotting. d) Schematic of identified MYPT1 O-GlcNAc sites, as well as the two deactivating phosphorylation sites and regions of MYPT1 responsible for critical protein-protein interactions. e) Endogenous MYPT1 is heavily and dynamically O-GlcNAc modified. NIH3T3 cells were treated with either 5SGlcNAc (200 μM), Thiamet-G (10 μM), or DMSO vehicle for 16, 20, or 20 h respectively. The O-GlcNAc modified proteins were then subjected to chemoenzymatic modification and then “mass-shifted” by PEGylation causing the modified fraction of proteins to run at higher molecular weights when analyzed by Western blotting. f-i) NIH3T3 cells stably expressing FLAG-tagged MYTP1 were treated under the indicated conditions before analysis by either mass-shifting or IP-Western blot. j) Schematic of the MYPT1Δ protein, which lacks the major O-GlcNAc region of the protein. k&l) MYPT1Δ loses a notable amount of O-GlcNAc. The O-GlcNAc levels of FLAG-tagged MYPT1 or MYPT1Δ were analyzed using mass-shifting or IP-Western blot.

Consistent with the high levels of modification we observed in our pulldown, recent proteomics experiments using the same chemoenzymatic strategy have identified many different endogenous MYPT1 O-GlcNAc sites^49–51^ localized around two different regions of the protein (Figure 4d). The most N-terminal of these areas, with modification at Ser379 and Thr381, is located near the portion of MYPT1 responsible for binding the phosphatase catalytic subunit PP1cδ and its substrate, phosphorylated MLC^52–54^. The second region contains eight identified O-GlcNAcylation sites (Ser566, Thr570, Thr577, Ser585, Ser589, Thr590, Thr592 & Thr594) clustered within a serine/threonine rich region, as well as ninth site slightly more distant at Thr637. These modification sites are located nearer to the inhibitory phosphorylation sites, and the portions of MYPT1 responsible for interacting with ROCK1/2 and the myosin motor complex^40,52–54^. Despite the clear potential for MYPT1 to be heavily O-GlcNAc modified, the exact stoichiometry of the endogenous modification had not been previously measured to our knowledge. To accomplish this, we again took advantage of the chemoenzymatic methodology (Figure 4a), but instead of installing an enrichment tag, we modified each O-GlcNAc moiety with a polyethylene glycol (PEG) chain of either 2 or 5 kDa in molecular weight. This mass-shifting approach causes the O-GlcNAc modified fraction of a protein to run higher on an SDS-PAGE gel for subsequent analysis by Western blotting^20,55^. We applied this technique to NIH3T3 cells under three sets of conditions: 5SGlcNAc treatment (200 μM) to lower O-GlcNAcylation levels, DMSO vehicle to leave them unchanged, or Thiamet-G treatment (10 μM) to raise them. We then visualized endogenous MYTP1 by Western blotting (Figure 4e). Under basal O-GlcNAc levels (i.e., DMSO treatment), we found that MYTP1 is indeed highly modified with essentially 100% of the protein running at mass-shifted molecular weights, and these modifications are largely removed upon treatment with 5SGlcNAc. With Thiamet-G treatment, we observed some potential further upward shift of the MYPT1 bands, suggesting an increase in O-GlcNAc levels. However, the most obvious change was an overall decrease in our ability to detect MYPT1. We believe that this is due to the reduced affinity of some antibodies to highly PEGylated proteins, which we previously found for Nup62^20^. In an attempt to overcome this issue, we next generated NIH3T3 cells that stably express a FLAG-tagged version of MYPT1 (Supplementary Figure 8a) and performed the same mass-shifting analysis with an anti-FLAG antibody (Figure 4f). Again, we saw near total O-GlcNAc modification of MYPT1 at multiple sites, and we confirmed that the modification stoichiometry could be dialed down or up upon 5SGlcNAc or Thiamet-G treatment, respectively. Unfortunately, we still detected some loss of anti-FLAG signal upon Thiamet-G treatment. Therefore, we next used an anti-FLAG immunoprecipitation (IP) to enrich MYPT1. We then observed the relative O-GlcNAc levels using an anti-O-GlcNAc antibody and confirmed that the protein’s modification status can be both down- and upregulated by inhibitor treatment (Figure 4g). Finally, we used both mass-shifting and anti-FLAG IP to demonstrate that MYTP1 O-GlcNAcylation levels will change in response to alterations in glucose concentration (1 vs. 9 g L^−1^) in the media (Figures 4h & i). Notably, these glucose-responsive differences in O-GlcNAc were smaller than those created by inhibitor treatment, in agreement with their more subtle effect on the cellular contraction phenotype (Figure 2e & f).

### O-GlcNAc of the MYTP1 serine/threonine rich region inhibits its phosphorylation and controls the sensitivity of cells to S1P-mediated contraction signaling

Previous experiments aimed at characterizing the MYPT1/ROCK interaction using recombinant proteins demonstrated that the serine/threonine region slightly inhibited this binding event^40^. Therefore, we hypothesized that O-GlcNAcylation in this location would increase this inhibition and prevent MYPT1 phosphorylation and inactivation. To test this possibility, we generated a series of mutant MYPT1 constructs (Supplementary Figure 9a). In order to generate these cells, we used human MYPT1, which is 93% identical to the mouse protein and enabled us to use RNA interference to selectively knockdown the endogenous MYPT1 in NIH3T3 cells (Supplementary Figure 8b). The first MYPT1 mutant, MYPT1(S/TtoA), contained serine and threonine mutations at each of the 8 sites in this region previously identified as O-GlcNAc modified by mass spectrometry^49–51^. Because OGT has been found to modify multiple residues in serine/threonine rich regions that cannot be readily differentiated by mass spectrometry, we also generated three deletion mutants: MYPT1(d564-578) and MYPT1(d588-602), which each lack one of the two stretches of serine/threonine, and MYPT1Δ where the entire serine/threonine-rich region (residues 550-600) was deleted. Importantly, all of these mutants displayed similar expression to stably expressed MYPT1 (Supplementary Figure 8a). With these cell lines in hand, we next used RNAi to knock down the endogenous copy of MYPT1, treated them with S1P (50 or 100 nM), and measured cellular contraction (Supplementary Figure 9b). At these S1P concentrations, we observed essentially no contraction in MYPT1, MYPT1(S/TtoA), or MYPT1(d564-578) expressing cells. In contrast, we found MYPT1Δ cells to have significant levels of contraction at both concentrations of S1P and MYPT1(d588-602) to have an intermediate phenotype. Consistent with a broad distribution of O-GlcNAc across the serine/threonine rich region of MYPT1, these results suggested that the entire domain contains modifications that are important for controlling the contraction phenotype. We then used the mass-shifting assay and IP-Western blotting to examine the O-GlcNAcylation stoichiometry of MYPT1Δ and found the levels noticeably but not completely reduced (Figure 4k & l), consistent with the localization of a reasonable fraction of O-GlcNAc to this region.

Next, we subjected MYPT1- or MYPT1Δ-expressing cells to the full range of S1P concentrations (0.05 - 5 μM) and measured the relative amounts of cell contraction (Figure 5a & b). We found that MYPT1Δ expressing cells were significantly more sensitive to S1P-mediated contraction signaling. Again, this phenotype was observed in multiple biological replicates (Supplementary Figure 10). To more directly test this model, we visualized the phosphorylation status of both MLC and MYPT1Δ after different lengths of treatment time with a low concentration of S1P (100 nM) (Figure 5c). As predicted, we observed dramatically less phosphorylation of both proteins in the cells expressing MYPT1 versus those expressing MYPT1Δ. These results are consistent with a model where O-GlcNAc in the serine/threonine region prevents MYPT1 phosphorylation at low concentrations of S1P and this regulatory system is lost in the MYTP1Δ mutant (Figure 5d).

**Figure 5.**
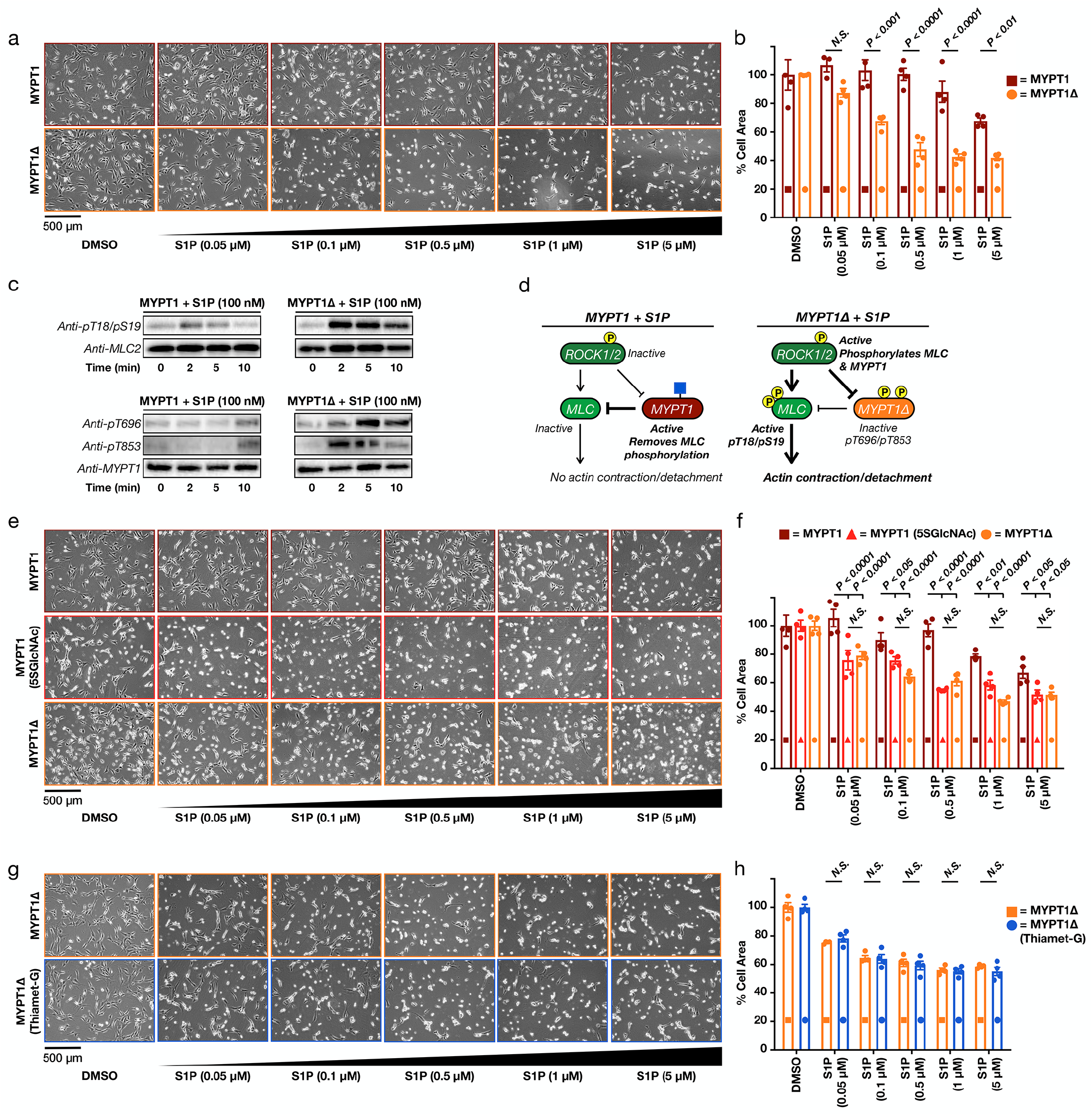
MYPT1 O-GlcNAc modification prevents its phosphorylation and deactivation, thereby inhibiting S1P-mediated contraction. a) MYTP1Δ expression sensitizes cells to S1P-mediated contraction. NIH3T3 cells stably expressing either MYPT1 or MYTP1Δ were transfected with RNAi to downregulate endogenous MYTP1. They were then treated with the indicated concentrations of S1P for 30 min. The contraction phenotype was visualized using bright-field microscopy. b) Quantitation of the data in (a). Results are the mean ± SEM of the relative culture plate area taken-up by cells in four randomly selected frames. Statistical significance was determined using a 2-way ANOVA test followed by Sidak’s multiple comparisons test. c) MYPT1Δ is more readily phosphorylated upon low S1P signaling. NIH3T3 cells stably expressing either MYPT1 or MYTP1Δ were transfected with RNAi to downregulate endogenous MYTP1. They were then treated with S1P (100 nM) for the indicated lengths of time. MLC and MYPT1/MYPT1Δ phosphorylation were analyzed by Western blotting. d) Our model. When MYPT1 is O-GlcNAc modified it is more resistant to phosphorylation and deactivation, thus maintaining the brakes on contraction. When MYTP1 O-GlcNAc is lost, it is phosphorylated, resulting in MLC phosphorylation and contraction. e) Direct MYPT1 O-GlcNAc modification is largely responsible for the phenotype. NIH3T3 cells stably expressing either MYPT1 or MYTP1Δ were transfected with RNAi to downregulate endogenous MYTP1. MYTP1-expressing cells were then treated with either DMSO or 5SGlcNAc (200 μM) for 16 h. Cells under all three sets of conditions were then treated with the indicated concentrations of S1P for 30 min. The contraction phenotype was visualized using bright-field microscopy. f) Quantitation of the data in (e). Results are the mean ± SEM of the relative culture plate area taken-up by cells in four randomly selected frames. Statistical significance was determined using a 2-way ANOVA test followed by Sidak’s multiple comparisons test. g) NIH3T3 cells stably expressing MYTP1Δ were transfected with RNAi to downregulate endogenous MYTP1. MYTP1Δ-expressing cells were then treated with either DMSO or Thiamet-G (10 μM) for 20 h. Cells were then treated with the indicated concentrations of S1P for 30 min. The contraction phenotype was visualized using bright-field microscopy. h) Quantitation of the data in (g). Results are the mean ± SEM of the relative culture plate area taken-up by cells in four randomly selected frames. Statistical significance was determined using a 2-way ANOVA test followed by Sidak’s multiple comparisons test.

Deletion of residues 550-600 did not compromise MYTP1Δ expression, limited O-GlcNAc modification, and resulted in the largest increased sensitivity to S1P. However, we were concerned that removal of these 50 amino acids could affect the functions of MYPT1. For example, MYPT1 can form a complex with OGT and direct it to protein substrates^47^. Therefore, we first examined whether MYPT1Δ expression altered overall O-GlcNAc by Western blotting. Importantly, we found no apparent changes in O-GlcNAc levels or OGT expression compared to MYPT1 (Supplementary Figure 8c). Second, we tested whether MYPT1Δ was still an active phosphatase. As above, we subjected MYPT1Δ cells to RNAi to remove endogenous MYPT1 and subjected the cells to S1P (100 nM) for a long time course of 180 min. We then examined the phosphorylation of MLC after 60 and 180 min. Consistent with increased S1P signaling upon expression of MYPT1Δ, we observed phosphorylation of MLC at 60 min, but this modification was removed by 180 min (Supplementary Figure 11a), demonstrating that MYPT1Δ can still dephosphorylate MLC to turn off the S1P-signaling pathway. Consistent with this biochemical result, we also observed the cells return to a relaxed state on the culture plate after 120 min (Supplementary Figure 11b). Together these data demonstrate that MYPT1Δ is still an active phosphatase despite deletion of the serine/threonine rich region.

Next, we set out to identify the mechanism by which O-GlcNAc would prevent MYPT1 phosphorylation by ROCK. We reasoned that O-GlcNAc mostly likely inhibits the physical interaction between the two proteins. To test this possibility, we treated NIH3T3 cells expressing FLAG-tagged MYPT1 with either DMSO or 5SGlcNAc (200 μM) and performed an anti-FLAG co-immunoprecipitation using the Catch and Release system (Thermo). We then blotted against endogenous ROCK, normalized the level to overall protein enrichment by Coomassie staining, and found that loss of O-GlcNAc resulted in a significant increase (~2-times) in the amount of ROCK interacting with MYPT1 (Supplementary Figure 12). These results support a model where O-GlcNAc blocks the protein-protein interaction between MYPT1 and ROCK.

Finally, we wanted to rule out the possibility that deletion of the serine/threonine region of MYPT1 has an effect beyond inhibiting O-GlcNAc and/or that O-GlcNAc on another protein contributes to the observed contraction. If neither of these possibilities is true, we would predict that cells expressing MYPT1Δ would respond to S1P in the same way as cells expressing wild-type MYPT1 treated with 5SGlcNAc. Accordingly, we compared three different conditions: cells stably expressing MYPT1, cells stably expressing MYPT1 in the presence of 5SGlcNAc, and cells stably expressing MYPT1Δ. In all cases RNAi was used to remove the endogenous copy of MYPT1 from the NIH3T3 cells. Subsequent treatment of these populations of cells with S1P (0.05 - 5 μM) demonstrated that MYPT1Δ sensitized cells to the same extent as 5SGlcNAc treatment (Figure 5e & f). Importantly, these data were recapitulated in a biological replicate (Supplementary Figure 13a).

We also used the inverse experiment to confirm this result, where one would predict that increasing O-GlcNAc in MYPT1Δ cells would not rescue the contraction phenotype. Thus, we treated MYPT1Δ expressing cells with either DMSO or Thiamet-G (10 μM) before subjecting them to S1P (0.05 - 5 μM) and quantitation of cell contraction (Figure 5g & h). In line with our expectation, we found that Thiamet-G did not alter the amount of contraction in these cells. Again, this result was identical in a biological replicate (Supplementary Figure 13b). These data are consistent with our model that MYPT1 O-GlcNAcylation is directly responsible for regulating this signaling pathway and that this is the major modification event controlling this process.

### O-GlcNAc inhibits S1P-mediated stressed collagen-matrix contraction by human dermal fibroblasts

Wound healing involves several overlapping phases carried out by dermal cells, including fibroblasts^56,57^. Fibroblasts migrate and proliferate into the matrix formed by the fibrin clot, secrete extracellular matrix (ECM) components, and differentiate into myofibroblasts to generate granulation tissue. The fibroblasts also physically contract themselves and the ECM to facilitate wound closure and remodeling. Three-dimensional collagen matrices are established *in vitro* models to study fibroblast behavior in tissue-like environments^21,58^, and a stressed collagen-matrix serves as a model for wound contraction (Supplementary Figure 14). In this model, dermal fibroblasts are mixed with collagen, allowed to polymerize, and then cultured for 16 - 48 h. During this time, the cells form actin stress fibers and focal adhesions. Subsequent release of the matrix from the culture plate and addition of certain signaling molecules, including S1P^22^, results in matrix contraction that can be quantified. To determine if O-GlcNAc levels control this process, we first examined immortalized human dermal fibroblast contraction in 2D cultures, analogous to the NIH3T3 experiments above. Accordingly, we treated dermal fibroblasts with either DMSO or 5SGlcNAc (200 μM) (Supplementary Figure 15a) for 16 h followed by S1P (0.05 - 5 μM). Similar to the NIH3T3 cells, we observed increased sensitivity of fibroblasts with low O-GlcNAc levels to S1P (Figure 6a & b and Supplementary Figure 15b).

**Figure 6.**
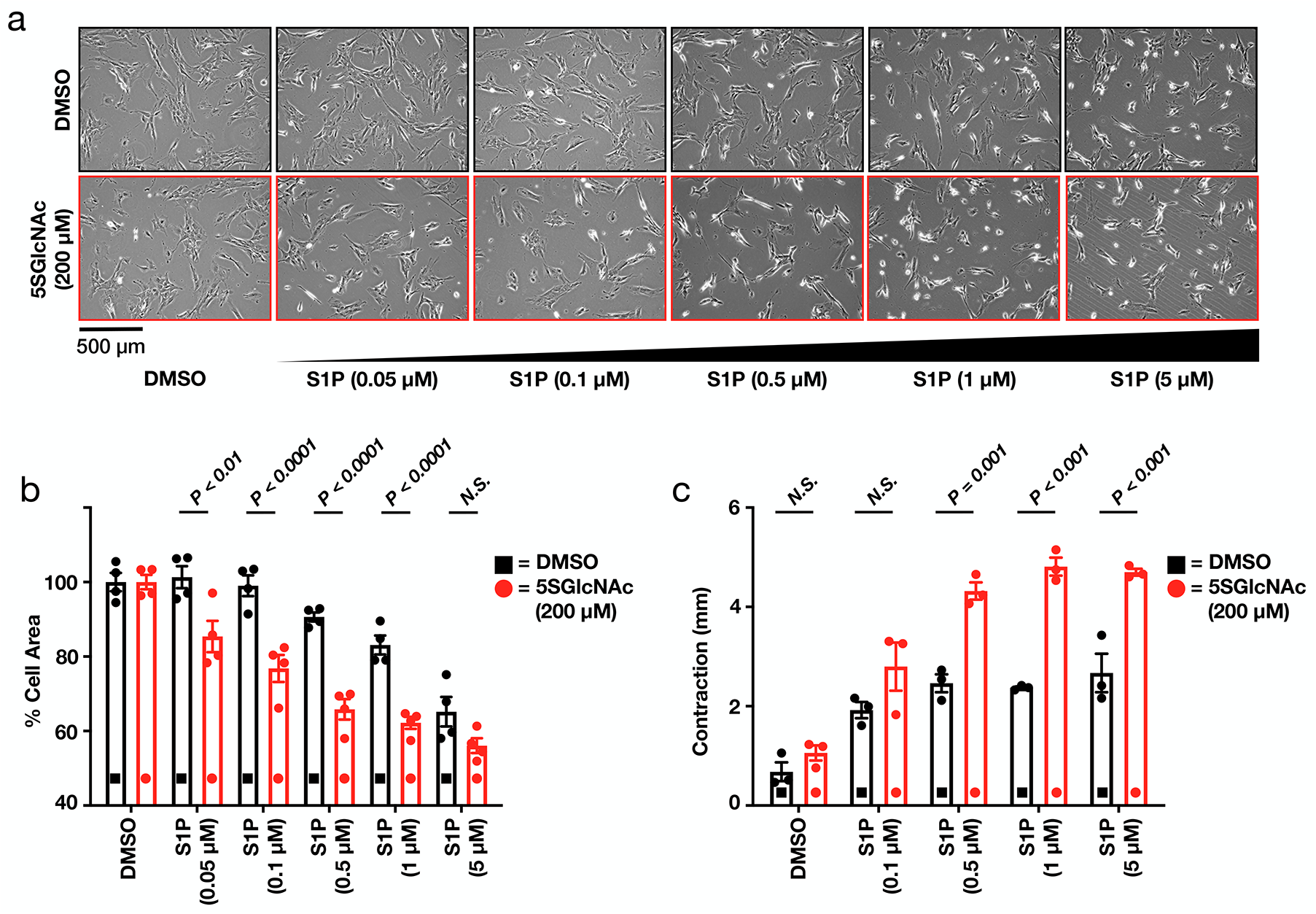
O-GlcNAc controls S1P-mediated collagen matrix contraction. a) Lowering O-GlcNAc levels increases the sensitivity of human dermal fibroblasts to S1P induced cell contraction in 2D culture. Fibroblasts were pre-treated with either DMSO or 5SGlcNAc (200 μM) for 16 h before addition of the indicated concentrations of S1P for 30 min. The contraction phenotype was then visualized using bright-field microscopy. b) Quantitation of the data in (a) Results are the mean ± SEM of the relative culture plate area taken-up by cells in four randomly selected frames. Statistical significance was determined using a 2-way ANOVA test followed by Sidak’s multiple comparisons test. c) Lowering O-GlcNAc increased the contraction of stressed collagen matrices in response to S1P. Contraction in millimeters was quantified as change in each matrix diameter from the initiation (t = 0 min) to termination (t = 30 min) of the assay. Results are the mean ± SEM of the millimeters contracted. Statistical significance was determined using a 2-way ANOVA test followed by Sidak’s multiple comparisons test.

Given this positive result, we then suspended dermal fibroblasts in collagen and plated them in 24-well dishes. After 30 h, the cells were treated with either DMSO or 5SGlcNAc (200 μM) and incubated for an additional 16 h. After this time, the fibroblasts were treated with a range of S1P concentrations (0.1 - 5 μM) and contraction was initiated by gently releasing the matrices from the culture plate. Images of the matrices were taken immediately (t = 0 min) and after 30 min (Supplementary Figure 16). The extent of contraction was then quantified by comparing the diameter of a matrix at 0 min to the diameter of the same matrix after 30 min (Figure 6c). Consistent with previous reports^22^, S1P caused matrix contraction by dermal fibroblasts with normal O-GlcNAc levels, but we observed significantly more contraction by cells treated with 5SGlcNAc. Together, these results demonstrate that O-GlcNAc levels control the contraction of human dermal fibroblasts in response to S1P and that this observation extends to a physiologically relevant model of wound healing.

## DISCUSSION

Taken together, these results strongly suggest that that MYPT1 O-GlcNAcylation functions as a nutrient sensor to regulate the sensitivity of fibroblasts to S1P-mediated contraction (Supplementary Figure 17). At this time, we do not yet know the biological consequences of this regulation in more complex setting, but there is reason to believe that they could be numerous. In the case of fibroblasts, S1P signaling has been shown to increase wound healing in mice^59,60^. We speculate that the elevated O-GlcNAc levels in diabetes may attenuate S1P signaling with detrimental consequences for fibroblast motility, differentiation, and contraction in wound healing. Importantly, our results in stressed-collagen matrices support this hypothesis, and we plan to continue this line of investigation. Furthermore, MYPT1 activity is critical for controlling smooth muscle contraction and therefore blood pressure. Several different GPCR agonists, including angiotensin, endothelin-1, S1P, epinephrine, and others activate the same Rho/ROCK pathway leading to MYPT1 phosphorylation and increased smooth muscle contraction^61–63^. Given our model where O-GlcNAcylation of MYTP1 acts downstream of ROCK, we hypothesize that this modification will control the sensitivity of smooth muscle cells to these various inputs, with potential implications in atherosclerosis and heart disease.

In summary, we present a biological model where O-GlcNAcylation of MYPT1 maintains its phosphatase activity by inhibiting the introduction of inactivating phosphorylation marks by ROCK1/2. This O-GlcNAcylation can then function in its well-established role as a sensor of the cellular environment and state to control the responsiveness of cells to contractile stimuli. In the past, the documented interaction between OGT and MYPT1 was thought to play only an adapter function for OGT substrate selection^47^, but we demonstrate here that O-GlcNAcylation of MYPT1 has its own important direct functions. Very recently, O-GlcNAc on MYPT1 was also shown to inhibit its phosphorylation by CDK1 with important consequences for the cell cycle^64^. We do not yet know the precise molecular mechanism by which O-GlcNAc inhibits MYPT1 phosphorylation. For example, we do not know if there are specific O-GlcNAc modification sites in the MYPT1 serine/threonine rich region that are most consequential, but our data suggests that our observations are likely a consequence of multiple O-GlcNAc modifications spread heterogeneously across the stretches of consecutive serine/threonine resides in this region. Thus, we believe that any individual site responsible for our observed phenotype probably does not exist. Our Co-IP suggests that O-GlcNAc inhibits the ROCK/ MYPT1 interaction. However, we cannot rule out the contribution of additional mechanisms. For example, O-GlcNAc might result in a conformation of MYPT1 that is refractory to phosphorylation in addition to having lower affinity for ROCK. Finally, our experiments comparing the effects of 5SGlcNAc or Thiamet-G treatment to MYTP1Δ expression cannot completely rule out some contribution of other O-GlcNAc modified proteins. Unfortunately, this “loss-of-modification” experiment is a general limitation of O-GlcNAc studies in cells, as to-date there are no robust methods for the site- or even protein-selective introduction of O-GlcNAcylation. However, we believe that the breadth of our experiments and heavy and dynamic nature of MYPT1 O-GlcNAc modification strongly support both our conclusions and the future exploration of these modifications in a variety of biological contexts.

## METHODS

Methods and any associated references are available in the online version of the paper.

## ACKNOWLEDGMENTS

The authors thank Prof. Frederick Grinnell at the UT Southwestern Medical Center for consultations on collagen matrix contraction. This research was supported by the American Cancer Society Research Scholar Grant (RSG-14-225-01-CCG), the University of Southern California, the Anton Burg Foundation, and the National Institutes of Health (R01GM125939) to M.R.P‥ N.J.P. is supported by NIGMS T32GM118289. The authors thank Prof. Kelley Moremen at the University of Georgia, who is supported by the National Institutes of Health (P41GM103390 and R01GM130915), for the generous gift of GalT(Y289L).

## AUTHOR CONTRIBUTIONS

N.J.P., A.R.B., N.D. and M.R.P. designed experiments and interpreted data. N.J.P. and A.R.B carried out cellular phenotype and Western blotting experiments. N.J.P. generated the MYPT1 stable cell lines and performed the associated experiments. N.J.P. also performed the co-IP, apoptosis, and collagen matrix experiments. N.D. performed the analysis of MYPT1 O-GlcNAc levels and dynamics. N.J.P, A.R.B. and M.R.P. prepared the manuscript.

## ADDITIONAL INFORMATION

Supplementary information is available in the online version of the paper. Reprints and permissions information is available online at http://www.nature.com/reprints/index.html. Correspondence and requests for materials should be addressed to M.R.P.

