## Supplemental Figures and Methods for "MYPT1 O-GlcNAc modification controls the sensitivity of fibroblasts to sphingosine-1-phosphate mediated cellular contraction"

### Table of contents:

**Supplementary Figure 1.** Modulating O-GlcNAcylation levels and identification of S1P. **Page S2**
**Supplementary Figure 2.** Lowering O-GlcNAc does not induce apoptosis. **Page S3**
**Supplementary Figure 3.** Cells return to a relaxed state after S1P-mediated contraction **Page S3**
**Supplementary Figure 4.** O-GlcNAc controls the sensitivity of fibroblasts to S1P mediated cell contraction (biological replicates of data in Figure 2). **Page S4**
**Supplementary Figure 5.** Schematic of the major S1P signaling pathways in mammalian cells **Page S5**
**Supplementary Figure 6.** O-GlcNAc controls signaling through the S1PR2 receptor. **Page S6**
**Supplementary Figure 7.** Inhibition of Rho kinase (ROCK1/2) blocks S1P-mediated cell contraction. **Page S7**
**Supplementary Figure 8.** Generation of MYPT1 stable cell-lines. **Page S7**
**Supplementary Figure 9.** Analysis of MYPT1 stable-cell lines **Page S8**
**Supplementary Figure 10.** MYTP1Δ expression sensitizes cells to S1P-mediated contraction (biological replicates of data in Figure 5). **Page S9**
**Supplementary Figure 11.** MYTP1Δ is an active phosphatase that can dephosphorylate MLC **Page S10**
**Supplementary Figure 12.** O-GlcNAc blocks the interaction between MYPT1 and ROCK **Page S11**
**Supplementary Figure 13.** Direct MYPT1 O-GlcNAcylation is largely responsible for the phenotype (biological replicate of data in Figure 5). **Page S12**
**Supplementary Figure 14.** Stressed-collagen matrix assay. **Page S12**
**Supplementary Figure 15.** O-GlcNAc controls S1P-mediated contraction of human dermal fibroblasts. **Page S13**
**Supplementary Figure 16.** Raw images fo the stressed collagen matrix assay. **Page S13**
**Supplementary Figure 17.** Our experimental model. **Page S14**
**Supplementary Figure 18.** Full blots from the corresponding Figures. **Page S15**
**Supplementary Figure 19.** Full blots from the corresponding Supplementary Figures. **Page S16**

**Experimental Methods** **Page S17**

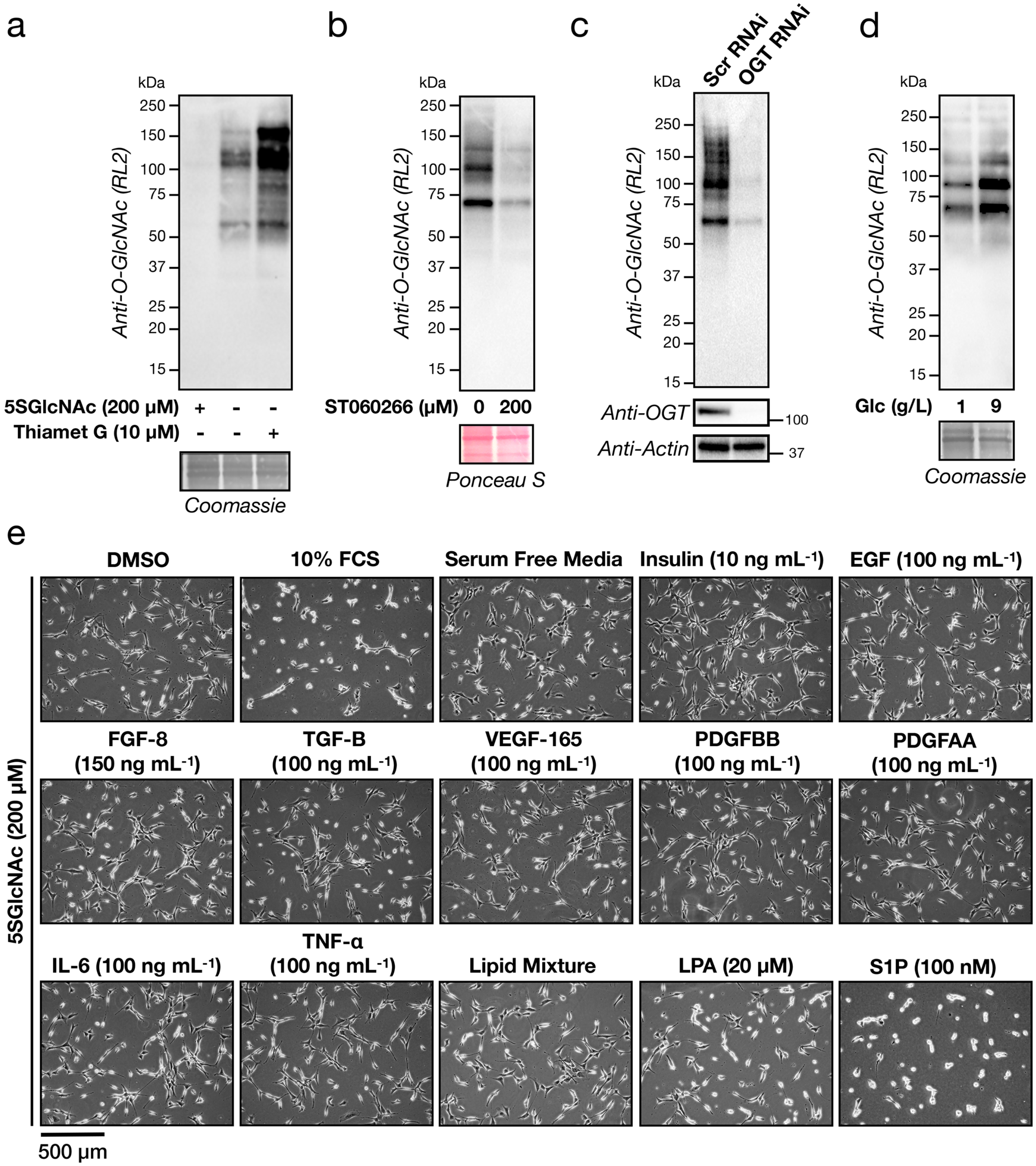


**Supplementary Figure 1. Modulating O-GlcNAcylation levels and identification of S1P.** a & b) O-GlcNAcylation levels can be changed upon OGT or OGA inhibition. NIH3T3 cells were treated with either the OGT inhibitors 5SGlcNAc (200 μM) or ST060266 (200 μM), Thiamet G (10 μM), or DMSO vehicle for 16, 20, or 20 h respectively before the O-GlcNAcylation levels were analyzed by Western blotting. b) O-GlcNAcylation levels can be lowered by OGT RNAi. NIH3T3 cells were transfected with RNAi targeting OGT or a scrambled sequence before analysis by Western blotting. d) O-GlcNAcylation levels can be changed by culturing cells in different glucose concentrations. NIH3T3 cells were cultured in the indicated concentrations of glucose for 48 h before analysis by Western blotting. e) Only serum- or S1P-treatment results in notable cell contraction when O-GlcNAcylation levels have been lowered. NIH3T3 cells were treated with DMSO or 5SGlcNAc (200 μM) for 16 h. The indicated signaling molecules were then added for 30 min, and the contraction phenotype was then visualized using bright-field microscopy.

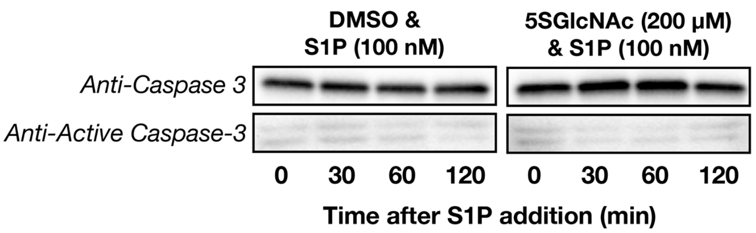


**Supplementary Figure 2. Lowering O-GlcNAc does not induce apoptosis.** NIH3T3 cells were treated with either DMSO or 5SGlcNAc (200 µM) for 16 h before addition of S1P (100 nM) for the indicated lengths of time. Activation of apoptosis was then ascertained by Western blotting against full-length and active caspase-3. No activation was observed under any of the experimental conditions.

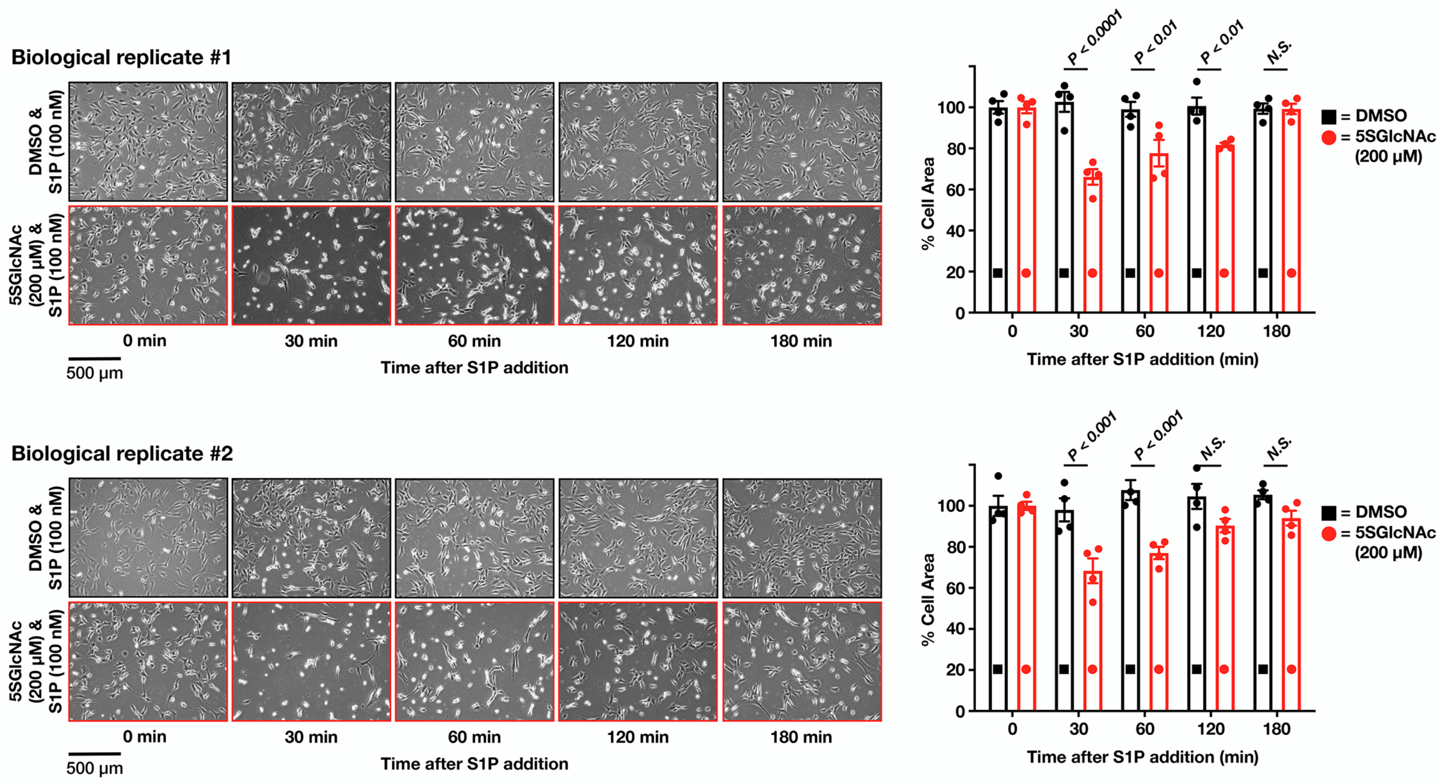


**Supplementary Figure 3. NIH3T3 cells return to a relaxed state after S1P-mediated contraction.** NIH3T3 cells that had been treated with either DMSO or 5SGlcNAc (200 µM) for 16 h before addition of the indicated concentrations of S1P (100 nM) for the indicated lengths of time. The contraction phenotype was then visualized using bright-field microscopy. Results are quantified as the mean ± SEM of the relative culture plate area taken-up by cells in four randomly selected frames. Statistical significance was determined using a 2-way ANOVA test followed by Sidak's multiple comparisons test.

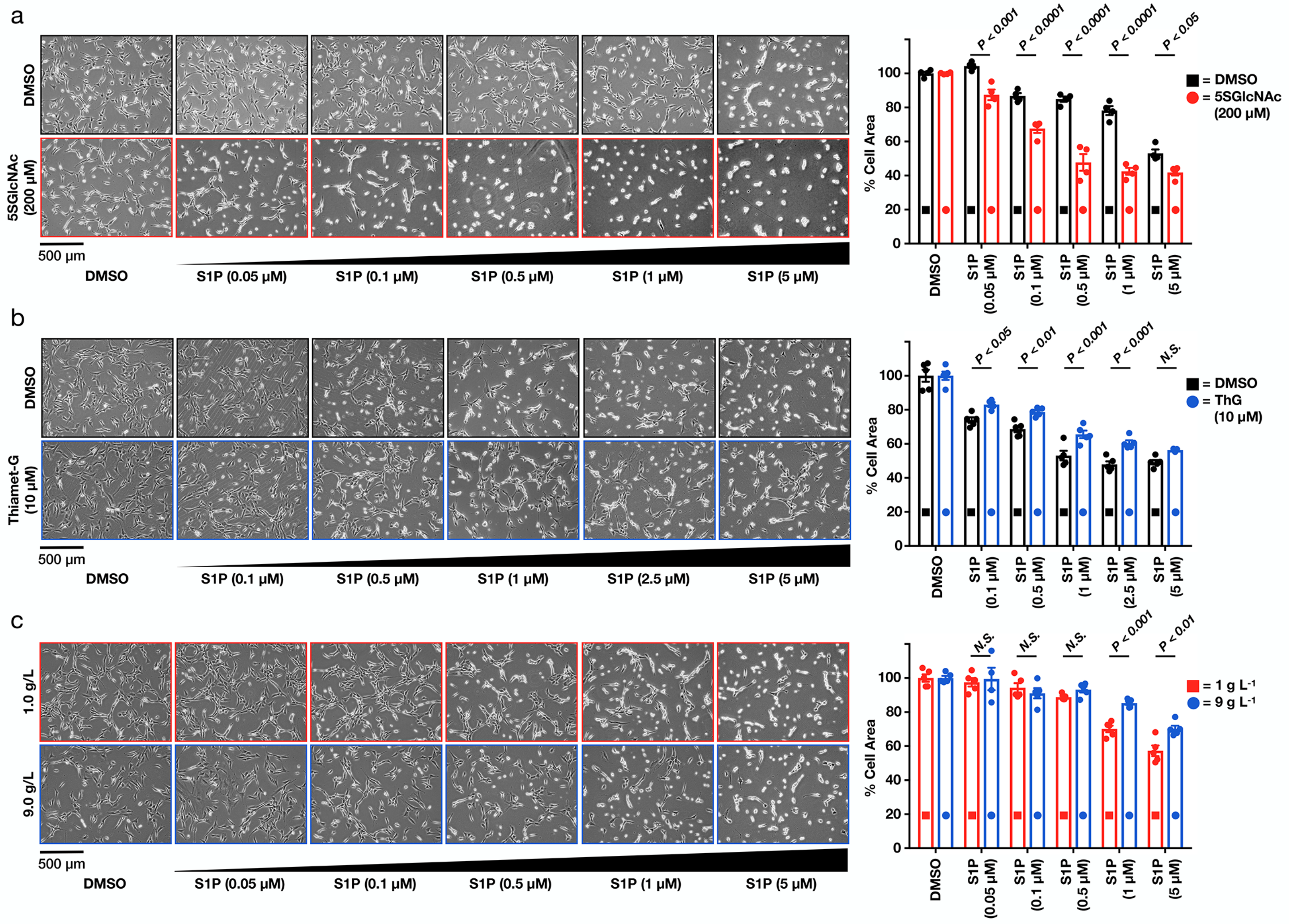


**Supplementary Figure 4. O-GlcNAc controls the sensitivity of fibroblasts to S1P mediated cell contraction (biological replicates of data in Figure 2).** a) Lowering O-GlcNAcylation levels increases the sensitivity of NIH3T3 cells to S1P induced cell contraction. NIH3T3 cells that had been treated with either DMSO or 5SGlcNAc (200 μM) for 16 h before addition of the indicated concentrations of S1P for 30 min. The contraction phenotype was then visualized using bright-field microscopy. Results are quantified as the mean ± SEM of the relative culture plate area taken-up by cells in four randomly selected frames. Statistical significance was determined using a 2-way ANOVA test followed by Sidak's multiple comparisons test. b) Raising O-GlcNAcylation levels decreases the sensitivity of NIH3T3 cells to S1P induced cell contraction. NIH3T3 cells that had been treated with either DMSO or the OGA inhibitor Thiamet-G (10 μM) for 20 h before addition of the indicated concentrations of S1P for 30 min. The contraction phenotype was then visualized using bright-field microscopy. Results are quantified as the mean ± SEM of the relative culture plate area taken-up by cells in four randomly selected frames. Statistical significance was determined using a 2-way ANOVA test followed by Sidak's multiple comparisons test. c) Glucose concentration controls the sensitivity of NIH3T3 cells to S1P induced cell contraction. NIH3T3 cells were cultured in the indicated amounts of glucose for 48 h before addition of the indicated concentrations of S1P for 30 min. The contraction phenotype was then visualized using bright-field microscopy. Results are quantified as the mean ± SEM of the relative culture plate area taken-up by cells in four randomly selected frames. Statistical significance was determined using a 2-way ANOVA test followed by Sidak's multiple comparisons test.

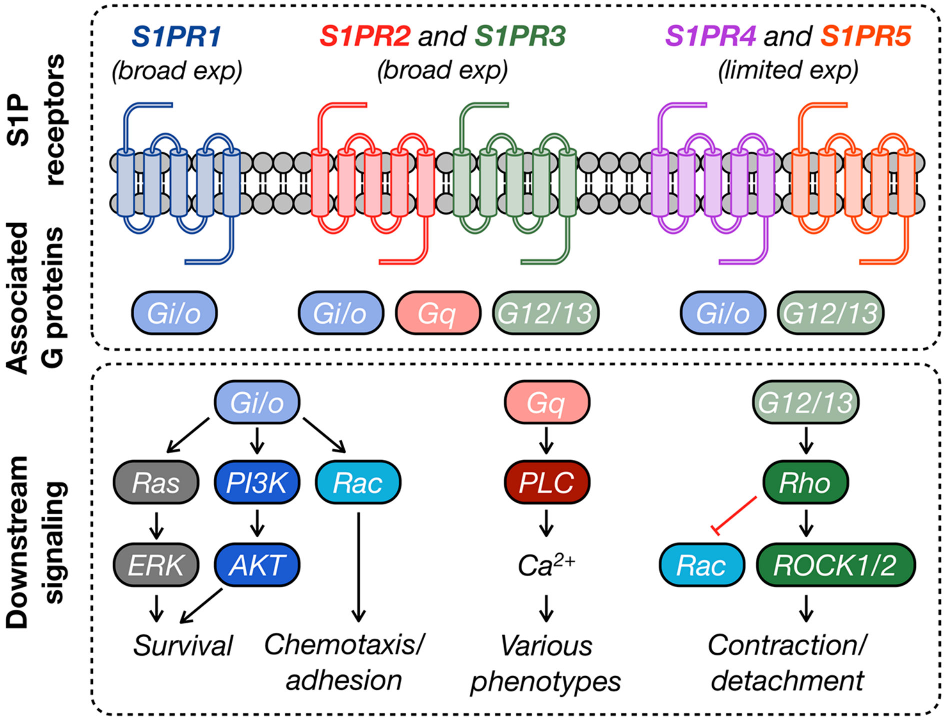


**Supplementary Figure 5. Schematic of the major S1P signaling pathways in mammalian cells.** S1P can agonize five different GPCRs (S1PR1 to R5) resulting in the potential activation of various downstream signaling pathways.

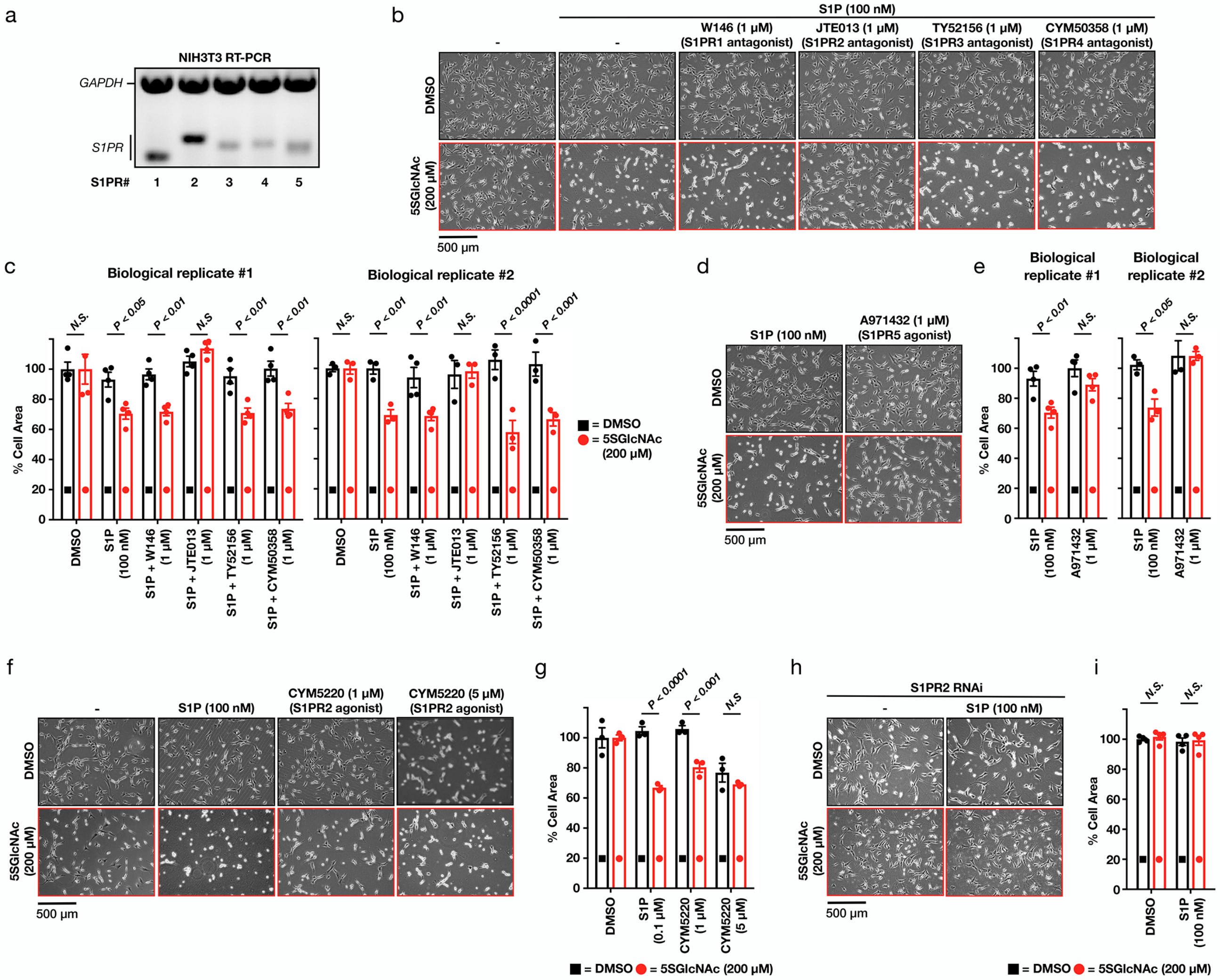


**Supplementary Figure 6. Signaling through the second S1P receptor, S1PR2, is responsible for the contraction phenotype.** a) NIH3T3 cells can express all five S1P GPCRs (S1PR1 to 5). mRNA was collected from NIH3T3 cells before being subjected to RT-PCR and visualization on an DNA-agarose gel. b) Antagonizing S1PR2, but not the other receptors, inhibits S1P-mediated cell contraction. NIH3T3 cells were treated with either DMSO or 5SGlcNAc (200 µM) for 16 h. The same cells were then treated with either additional DMSO or the indicated selective antagonists (1 µM) for 10 min followed by S1P (100 nM) for 30 min. The contraction phenotype was then visualized using bright-field microscopy. c) Quantitation of the data in (b). Results are the mean ± SEM of the relative culture plate area taken-up by cells in four randomly selected frames. Statistical significance was determined using a 2-tailed student's t-test. d) S1PR5 agonism does not induce cell contraction. NIH3T3 cells were treated with either DMSO or 5SGlcNAc (200 µM) for 16 h. The same cells were then treated with either S1P (100 nM) or the S1PR5-selective agonist A971432 (1 µM) for for 30 min. The contraction phenotype was then visualized using bright-field microscopy. e) Quantitation of the data in (d). Results are the mean ± SEM of the relative culture plate area taken-up by cells in four randomly selected frames. Statistical significance was determined using a 2-tailed student's t-test. f) Lowering O-GlcNAcylation levels increases the sensitivity of NIH3T3 cells to S1PR2 induced cell contraction. NIH3T3 cells that had been treated with either DMSO or 5SGlcNAc (200 µM) for 16 h before addition of the indicated concentrations of the S1PR2-selective agonist CYM5220 for 30 min. The contraction phenotype was then visualized using bright-field microscopy. g) Quantitation of the data in (f). Results are the mean ± SEM of the relative culture plate area taken-up by cells in four randomly selected frames. Statistical significance was determined using a 2-way ANOVA test followed by Sidak's multiple comparisons test. h) S1PR2 knockdown using siRNA blocks contraction phenotype. NIH3T3 cells were transfected with either scramble or S1Pr2-targeted RNAi for 48 h before addition of either more DMSO or S1P (100 nM) for 30 min. The contraction phenotype was then visualized using bright-field microscopy. i). Quantitation of the data in (h). Results are the mean ± SEM of the relative culture plate area taken-up by cells in four randomly selected frames. Statistical significance was determined using a 2-way ANOVA test followed by Sidak's multiple comparisons test.

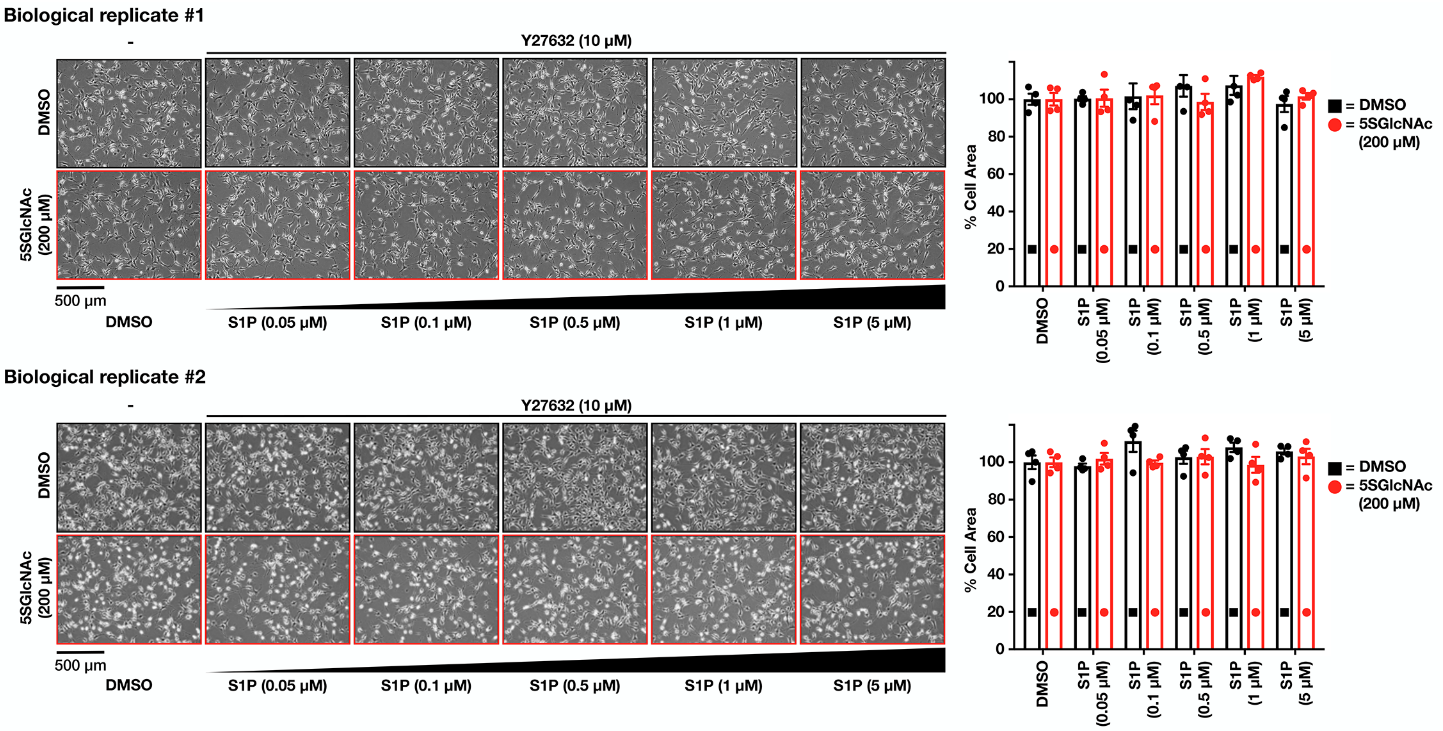


**Supplementary Figure 7. Inhibition of Rho kinase (ROCK1/2) blocks S1P-mediated cell contraction.** NIH3T3 cells were treated with either DMSO or 5SGlcNAc (200 µM) for 16 h. The same cells were then treated with either additional DMSO or the ROCK1/2 inhibitor Y27632 (10 µM) for 1 h followed by the indicated concentrations of S1P for 30 min. The contraction phenotype was then visualized using bright-field microscopy. The results were then quantified and are presented as mean ± SEM of the relative culture plate area taken-up by cells in four randomly selected frames.

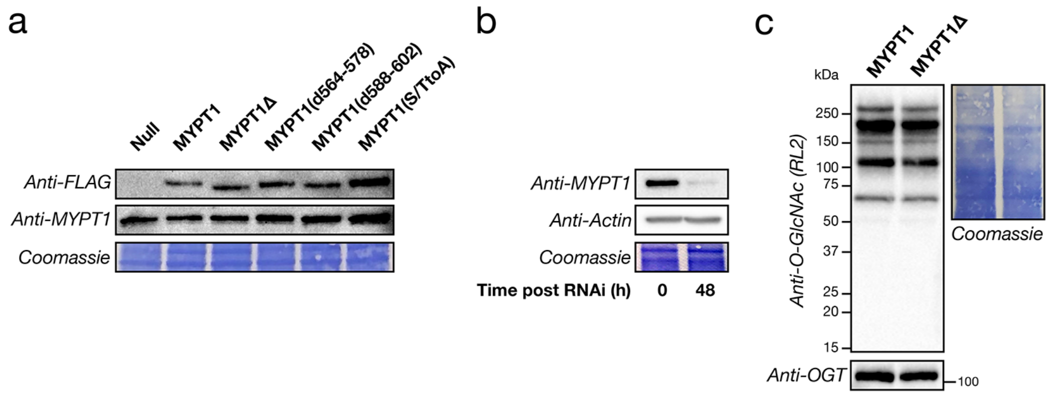


**Supplementary Figure 8. Characterization of MYPT1 stable cell-lines.** a) Generation of cells stably expressing MYPT1 or its associated mutants. NIH3T3 cells were stably transfected with either FLAG-tagged human MYTP1 or the indicated MYPT1 mutants using the PiggyBac retrotransposon system before analysis by Western blotting. b) RNAi efficiently knocks-down endogenous MYPT1. NIH3T3 cells were transfected with RNAi targeting mouse MYPT1 (Sigma) and protein levels were measured by Western blotting. c) MYPT1Δ expression does not adversely affect O-GlcNAc modification. O-GlcNAc modifications in MYPT1 and MYPT1Δ expressing cells were visualized by Western blotting.

a

```
           MYPT1: 550-HKSCSFGRRQDDLISSSVPSTTSTPTVTSAAGLQKSLLSSTSTTTKITTGSSSA-603
  MYPT1(S/TtoA): 550-HKSCSFGRRQDDLISSAVPSATSTPTVASAAGLQKALLSAASATAKITTGSSSA-603
MYPT1(d564-578): 550-HKSCSFGRRQDDLI---------------AAGLQKSLLSSTSTTTKITTGSSSA-603
MYPT1(d588-602): 550-HKSCSFGRRQDDLISSSVPSTTSTPTVTSAAGLQKSLL---------------A-603
          MYPT1Δ: 550-H----------------------------------------------------SSA-603
```

b

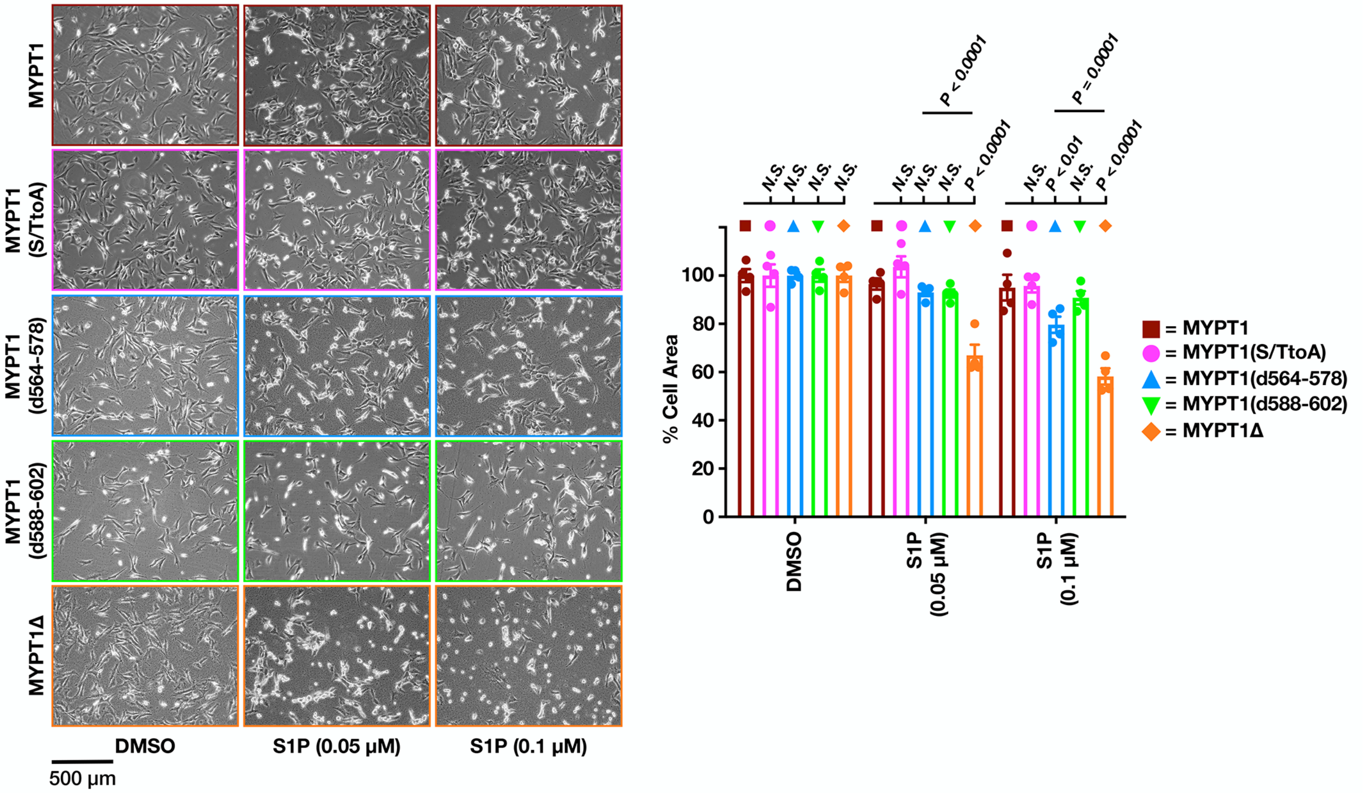


**Supplementary Figure 9. Analysis of MYPT1 mutants shows that deletion of the serine/threonine domain sensitizes cells to S1P-mediated contraction.** a) Sequence alignment of MYPT1 and the different mutants tested here. Red indicates potential O-GlcNAc modification sites (S or T), and blue indicates O-GlcNAc sites previously identified by mass spectrometry. MYTP1(S/TtoA) mutates all of the previously identified O-GlcNAc sites, MYPT1(d564-578) deletes the first serine/threonine rich region, MYPT1(d588-602) deletes the second serine/threonine rich region, and MYPT1Δ deletes the entire serine/threonine rich domain. b) MYPT1Δ, and to a lesser extent MYPT1(d564-578), sensitizes cells to S1P-mediated contraction. NIH3T3 cells expressing the indicated MYPT1 proteins and the endogenous copy was removed by RNAi. DMSO or S1P was then added and the contraction of the cells was measured after 30 min. Results are the mean ± SEM of the relative culture plate area taken-up by cells in four randomly selected frames. Statistical significance was determined using a 2-way ANOVA test followed by Sidak's multiple comparisons test.

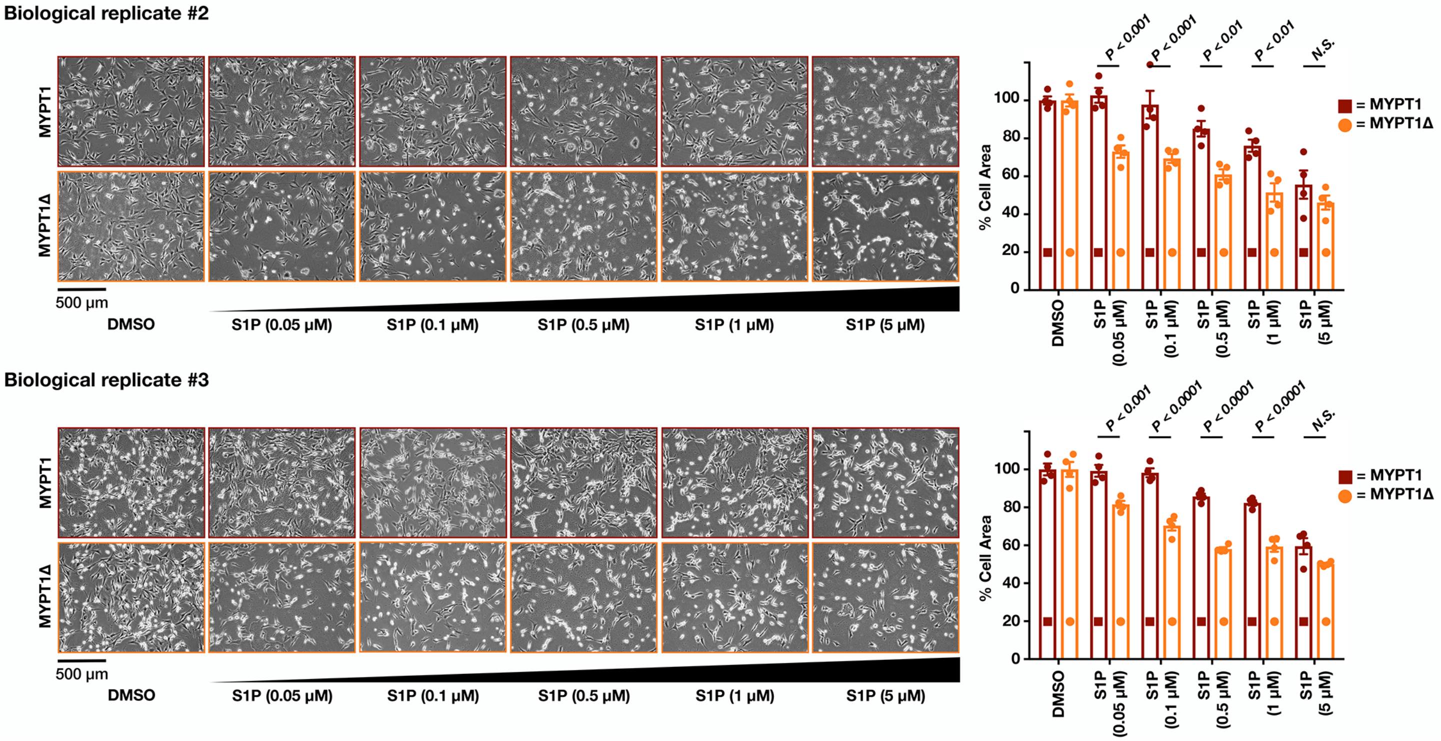


**Supplementary Figure 10. MYTP1Δ expression sensitizes cells to S1P-mediated contraction (biological replicates of data in Figure 5).** NIH3T3 cells stably expressing either MYPT1 or MYTP1Δ were transfected with RNAi to downregulate endogenous MYTP1. They were then treated with the indicated concentrations of S1P for 30 min. The contraction phenotype was visualized using bright-field microscopy. The results were then quantified and are presented as mean ± SEM of the relative culture plate area taken-up by cells in four randomly selected frames. Statistical significance was determined using a 2-way ANOVA test followed by Sidak's multiple comparisons test.

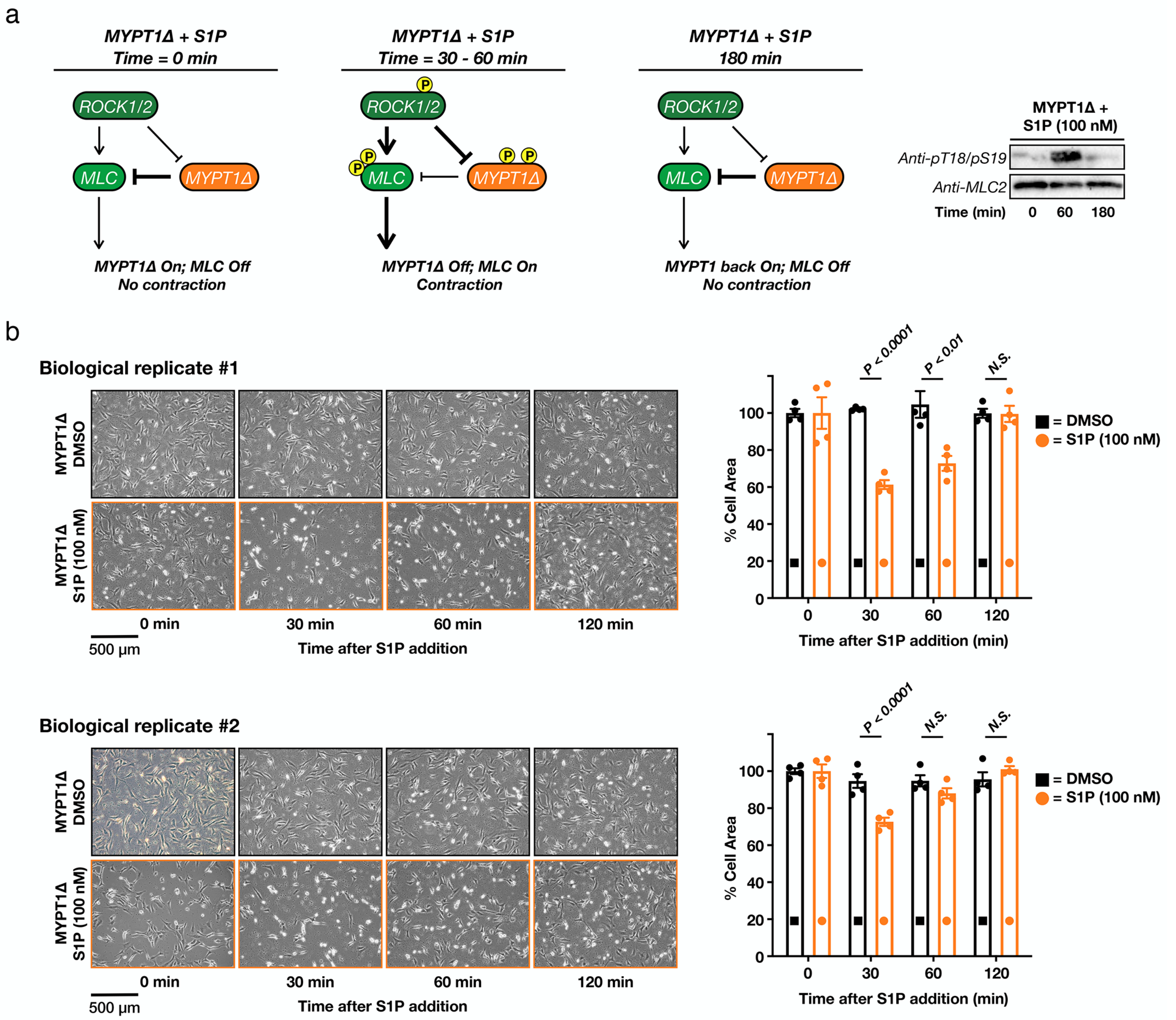


**Supplementary Figure 11. MYPT1Δ is an active phosphatase that can dephosphorylate MLC and return cells to a relaxed state.** a) If MYPT1Δ is an active phosphatase, we expect that it will become phosphorylated and deactivated by ROCK after S1P treatment but will return to a desphosphorylated and active state after a longer period of time. This will result in MLC dephosphorylation and relaxation of the cells. This is exactly what we observed by Western blotting. b) Cells expressing MYPT1Δ return to a relaxed state over 180 min. The contraction phenotype was visualized using bright-field microscopy. The results were then quantified and are presented as mean ± SEM of the relative culture plate area taken-up by cells in four randomly selected frames. Statistical significance was determined using a 2-way ANOVA test followed by Sidak's multiple comparisons test.

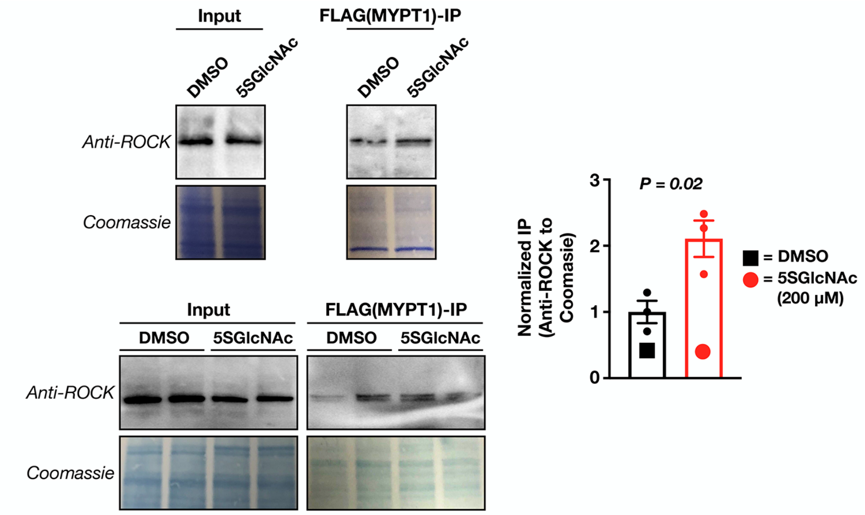


**Supplementary Figure 12. O-GlcNAc blocks the interaction between MYPT1 and ROCK.** NIH3T3 cells expressing flag-tagged MYPT1 were treated with either DMSO or 5SGlcNAc (200 µM). An anti-flag co-immunoprecipitation was then performed using the Catch and Release system (Thermo). ROCK enrichment was detected by Western blotting and normalized to overall protein capture (Coomassie staining). The results were quantities and presented as mean ± SEM of the normalized ROCK levels (n = 3 biological replicates). Statistical significance was determined using a 2-tailed, unpaired Student's t-test.

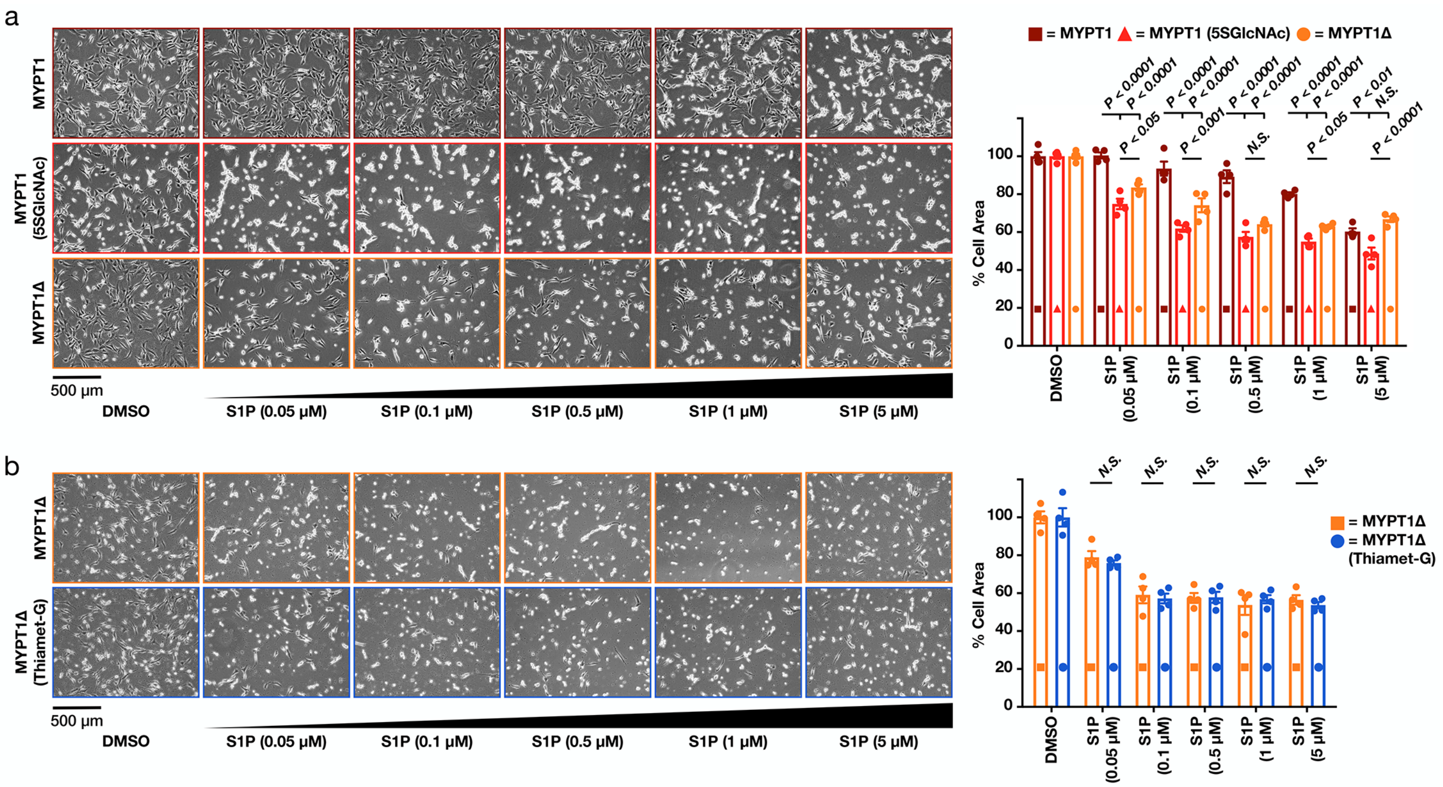


**Supplementary Figure 13. Direct MYPT1 O-GlcNAcylation is largely responsible for the phenotype (biological replicates of data in Figure 5).** a) NIH3T3 cells stably expressing either MYPT1 or MYTP1Δ were transfected with RNAi to downregulate endogenous MYTP1. MYTP1-expressing cells were then treated with either DMSO or 5SGlcNAc (200 µM) for 16 h. Cells under all three sets of conditions were then treated with the indicated concentrations of S1P for 30 min. The contraction phenotype was visualized using bright-field microscopy. The results were then quantified and are presented as mean ± SEM of the relative culture plate area taken-up by cells in four randomly selected frames. Statistical significance was determined using a 2-way ANOVA test followed by Sidak's multiple comparisons test. b) NIH3T3 cells stably expressing either MYPT1 or MYTP1Δ were transfected with RNAi to downregulate endogenous MYTP1. MYTP1-expressing cells were then treated with either DMSO or Thiamet-G (10 µM) for 20 h. Cells under all three sets of conditions were then treated with the indicated concentrations of S1P for 30 min. The contraction phenotype was visualized using bright-field microscopy. The results were then quantified and are presented as mean ± SEM of the relative culture plate area taken-up by cells in four randomly selected frames. Statistical significance was determined using a 2-way ANOVA test followed by Sidak's multiple comparisons test.

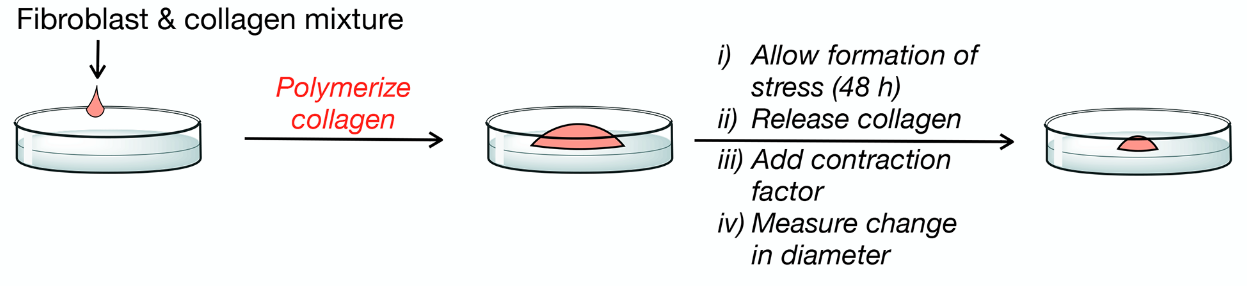


**Supplementary Figure 14. Stressed-collagen matrix assay.** Mixing fibroblasts in a collagen matrix generates a 3D culture model of tissue that allows fibroblast contraction (stressed matrix) to be measured in response to S1P.

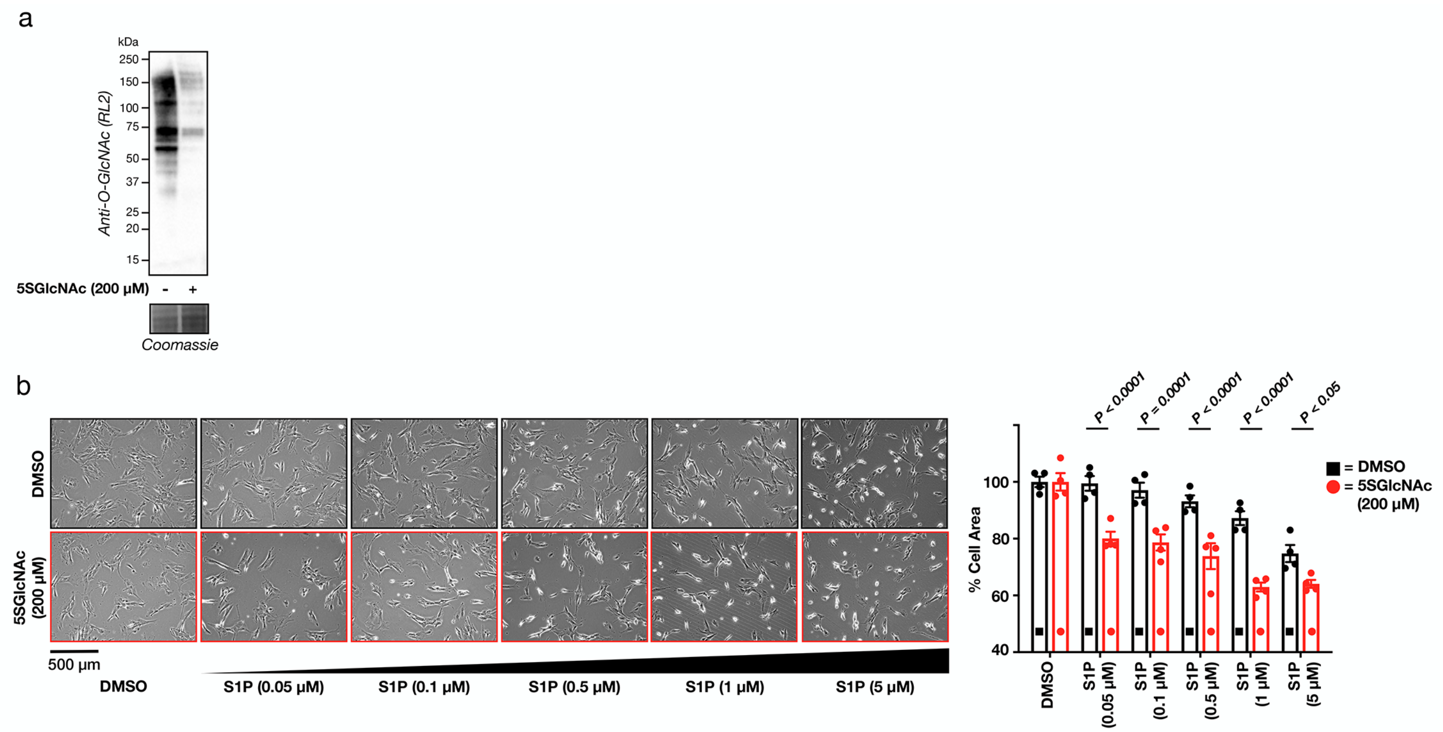


**Supplementary Figure 15. O-GlcNAc controls S1P-mediated contraction of human dermal fibroblasts.** a) O-GlcNAcylation levels can lowered upon OGT inhibition. Primary human dermal fibroblasts were treated with either the OGT inhibitor 5SGlcNAc (200 µM) for 16 h before the O-GlcNAcylation levels were analyzed by Western blotting. b) Lowering O-GlcNAc levels increases the sensitivity of human dermal fibroblasts to S1P induced cell contraction in 2D culture (biological replicate of Figure 6a & b). Fibroblasts were pre-treated with either DMSO or 5SGlcNAc (200 µM) for 16 h before addition of the indicated concentrations of S1P for 30 min. The contraction phenotype was visualized using bright-field microscopy. The results were then quantified and are presented as mean ± SEM of the relative culture plate area taken-up by cells in four randomly selected frames. Statistical significance was determined using a 2-way ANOVA test followed by Sidak's multiple comparisons test.

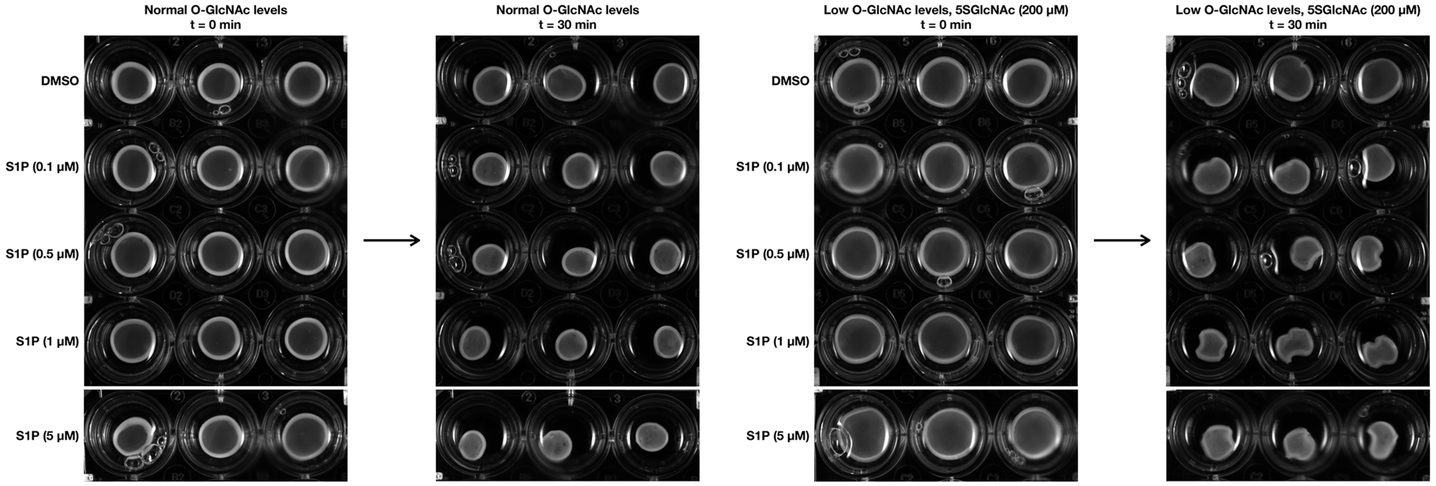


**Supplementary Figure 16. Raw images fo the stressed collagen matrix assay.** Stressed collagen matrices containing human dermal fibroblasts with either normal O-GlcNAc levels or low O-GlcNAc levels, 5SGlcNAc (200 µM), were treated with the indicated concentrations of S1P. The matrices were gently released and images were taken immediately (t = 0 min) and after 30 min.

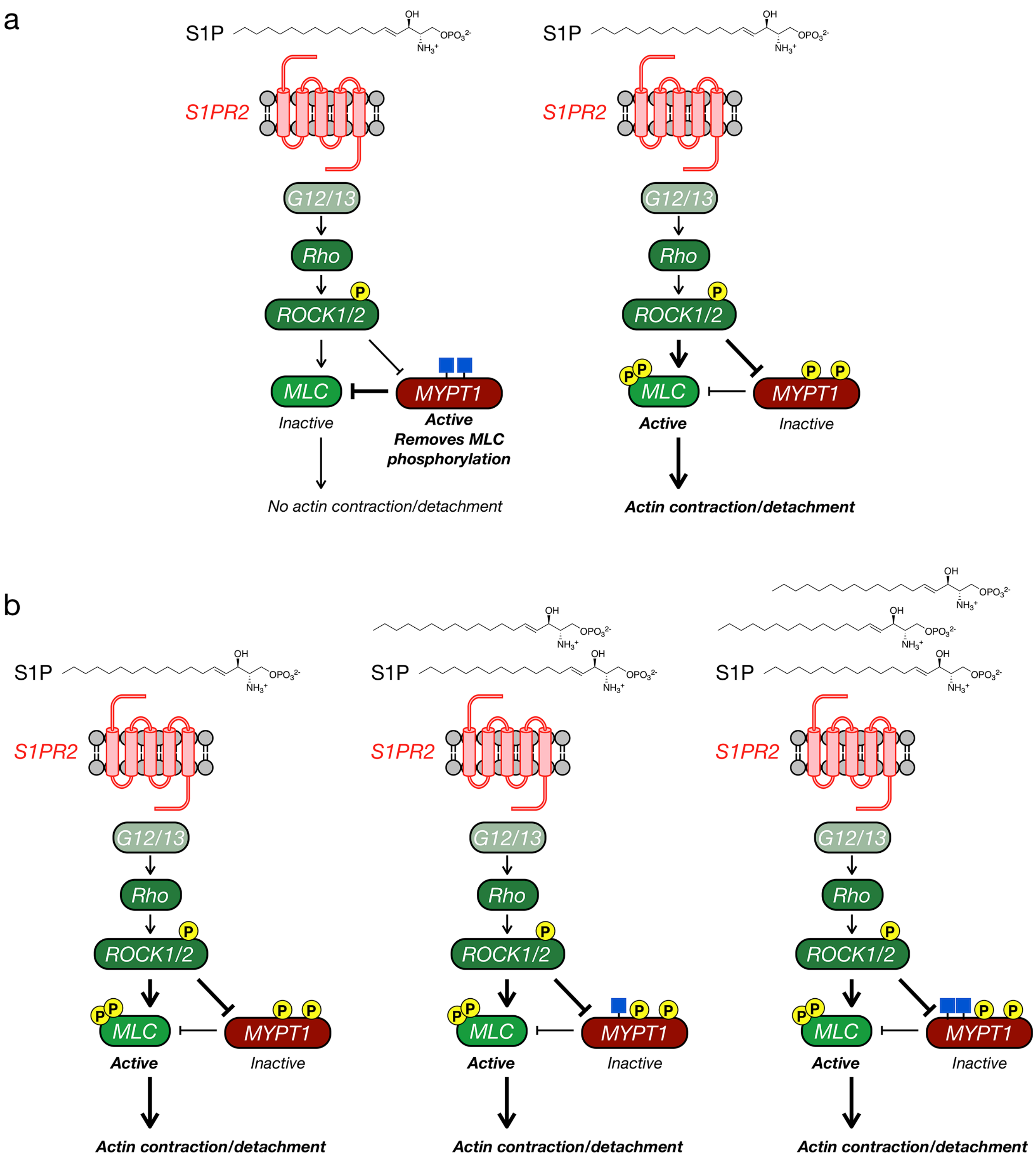


**Supplementary Figure 17. Our experimental model.** a) MYPT1 O-GlcNAcylation inhibits its phosphorylation by ROCK1/2. This maintains MYPT1 phosphatase activity, resulting in inactive MLC and no actin contraction. Loss of O-GlcNAc enables ROCK1/2 to phosphorylate and deactivate MYTP1. b) Therefore, MYPT1 O-GlcNAcylation levels control the sensitivity of cells to the concentration of S1P, where more MYTP1 O-GlcNAcylation requires more S1P to illicit actin contraction and cell detachment.

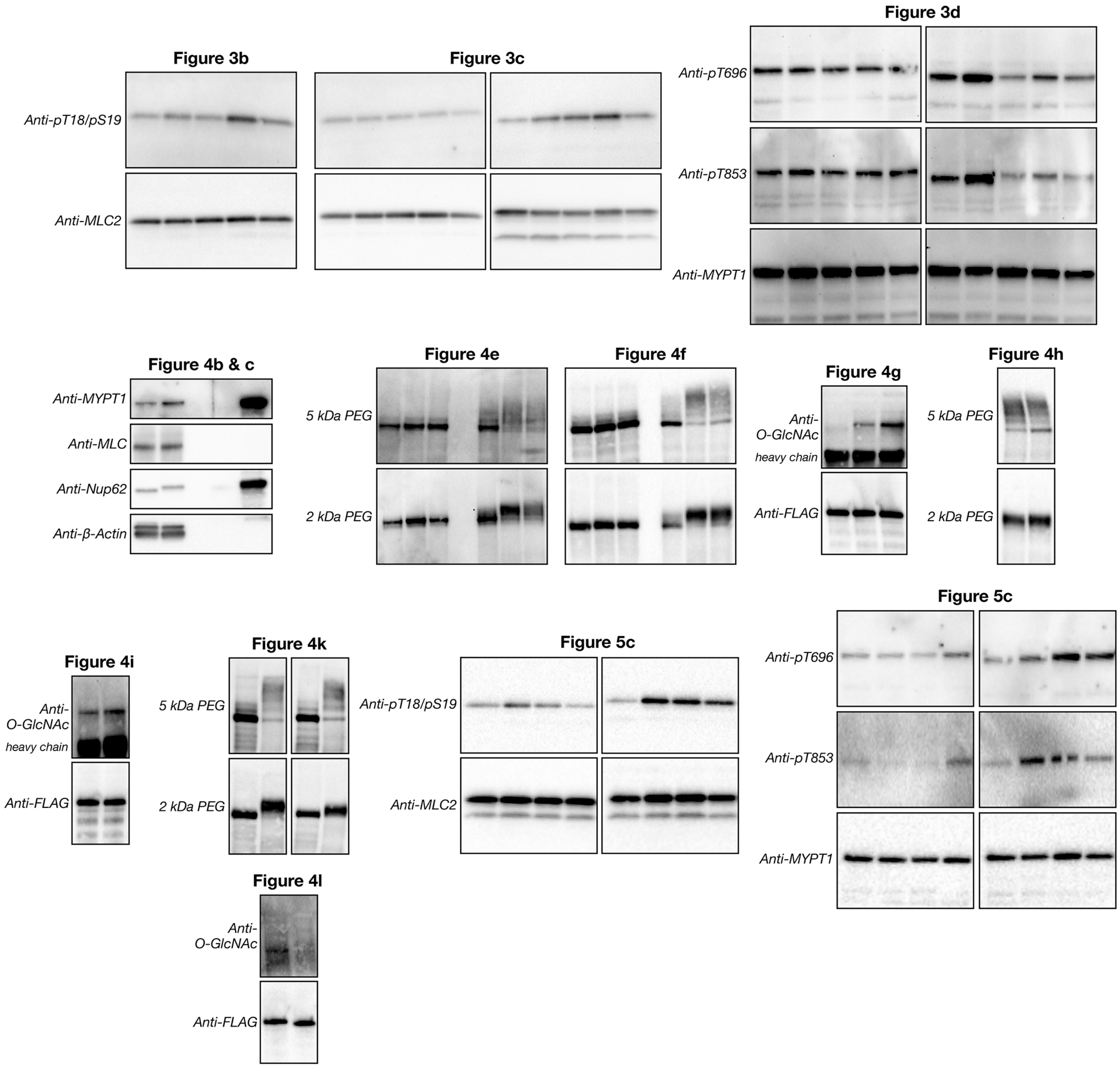


**Supplementary Figure 18. Full blots from the corresponding Figures.**

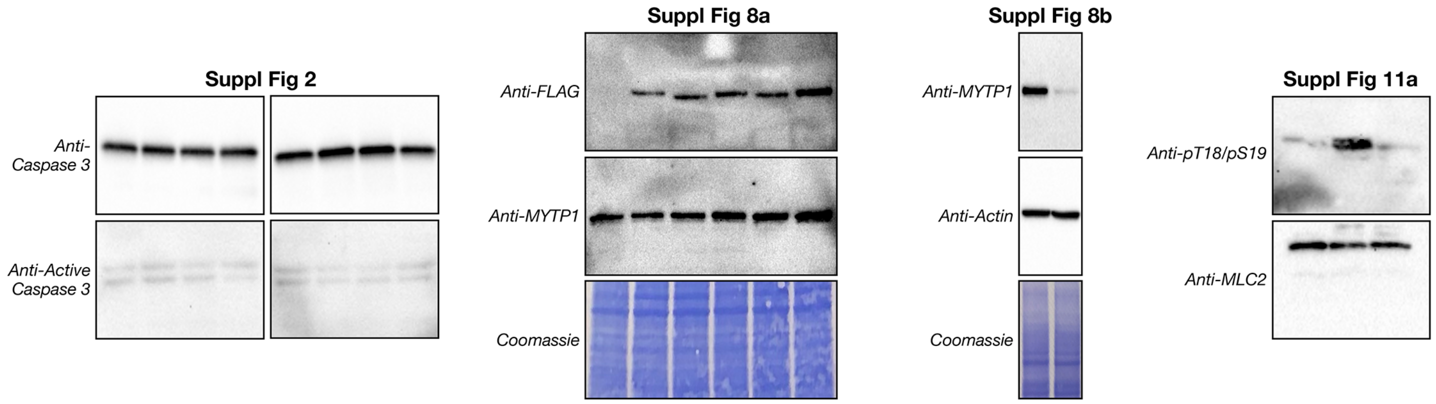


**Supplementary Figure 19. Full blots from the corresponding Supplementary Figures.**

### EXPERIMENTAL METHODS

#### Synthesis of known small molecules.

Known compounds Thiamet-G{Yuzwa:2008fq} and $Ac_45SGlcNAc${Gloster:2011da} were synthesized according to literature procedures. Both were dissolved as 1,000x stocks in DMSO.

#### Cell culture.

Mouse embryonic fibroblast cell-line NIH3T3 (ATCC) was propagated in DMEM medium (Genessee Scientific) supplemented with 10% calf serum (Atlanta Biologics). NIH3T3 cell-lines stably expressing either FLAG-tagged MYPT1, MYPT1(S/TtoA), MYPT1(d564-578), MYPT1(d588-602) or MYPT1Δ were grown in DMEM + 10% FCS supplemented with 250 ng $mL^{-1}$ puromycin (1 mg $mL^{-1}$ stock in $H_2O$). Human dermal fibroblast cell-line BJ-5ta (ATCC) was propagated in 4:1 DMEM:Medium 199 (ATCC) + 10% FBS supplemented with 0.01 mg $mL^{-1}$ hygromycin B (10 mg $mL^{-1}$ stock in $H_2O$). All cell-lines were grown in a humidified incubator at 37 °C and 5% $CO_2$ atmosphere.

#### Antibodies.

All antibodies were incubated in OneBlock™ Western-CL blocking buffer purchased from Genesee Scientific (20-313). Anti-FLAG-Tag (2368S), anti-MYPT1 (2634S), anti-MLC2 (3672S), anti-pT18/pS19 MLC (3674S), anti-pT696 MYPT1 (5163S), anti-pT853 MYPT1 (4563S), anti-OGT (24083S), anti-Caspase-3 (9762), and anti-Cleaved Caspase-3 (9664) were purchased from Cell Signaling Technology. Anti-Nup62 (610497) was purchased from BD Biosciences. Anti-RL2 (MA1-072) was purchased from ThermoFisher Scientific. Anti-β-actin (A5441) and Anti-Rock1/2 (07-1458) was purchased from MilliporeSigma. Horseradish peroxidase (HRP) conjugated secondary antibodies were purchased from Jackson ImmunoResearch.

#### General procedure for Western blotting.

Post lysis, proteins were separated by SDS-PAGE (200 V, 45 min) before being transferred to a PDVF membrane (Bio-Rad) using standard procedures. Western blots were blocked in OneBlock™ Western-CL blocking buffer (Genesee) for 1 h at rt. Blots were then incubated with primary antibody in fresh blocking buffer at 4 °C overnight. All primary antibodies were used at 1:1,000 dilution unless otherwise indicated. Following overnight incubation, blots were washed in TBST (Cell Signaling, 3x 10 min) and incubated with horse radish peroxidase (HRP) conjugated secondary antibody in fresh blocking buffer for 1 h at rt. All HRP-secondaries were used at 1:10,000 dilution. Once complete, blots were again washed in TBST (3x 10 min). Blots were developed using ECL reagents (Bio-Rad) and the ChemiDoc XRS+ molecular imager (Bio-Rad).

#### General procedure for 2D contraction assay.

For initial characterization of phenotype, NIH3T3 cells were seeded at 1 x $10^5$ cells in 6-well dishes 8 h prior to treatment with either DMSO vehicle, 5SGlcNAc (200 μM), or Thiamet-G (10 μM). Following incubation for 16 or 20 h for 5SGlcNAc or Thiamet-G/DMSO respectively, cells were treated with calf serum (Atlanta Biologics, 10% by volume) or fresh serum free media for 30 min. Each well was analyzed using a Leica Microscope to capture bright-field microscopy images at 20x magnification. Quantification of contraction phenotype was determined by taking the mean ± SEM of the relative culture plate area covered by cells in four randomly selected frames per well. Images were analyzed using Adobe Photoshop. Background pixels were selected using the Magic Wand tool and subtracted from total pixels. This value was then normalized using a control, untreated well allowing for quantification of the difference in area taken up by cells before and after contraction. Statistical significance was determined using a 2-way ANOVA test followed by Sidak's multiple comparisons test. Representative images for each treatment were selected from one of the four frames used for quantification.

#### Serum screen.

Seeded NIH3T3 cells were treated with 5SGlcNAc (200 μM) as stated above. After 16 h, wells were treated with one of the following; Fetal Calf Serum (10% v/v), replacement with serum free media, VEGF-165 (100 ng $mL^{-1}$), EGF (100 ng $mL^{-1}$), IGF (10 ng $mL^{-1}$), TGF-β (100 ng $mL^{-1}$), TNF-α (100 ng $mL^{-1}$), FGF-8 (150 ng $mL^{-1}$), PDGF-AA (100 ng $mL^{-1}$), PDGF-BB (100 ng $mL^{-1}$), IL-6 (100 ng $mL^{-1}$), LPA (20 μM), Lipid Mixture (MilliporeSigma, L0288-100ML), or S1P (100 nM). After a 30 min incubation at 37 °C, contraction for each treatment was measured as indicated above.

**OGT inhibition with ST060266.**
Seeded NIH3T3 cells were treated with covalent OGT inhibitor ST060266 (MilliporeSigma, 200 μM) for 16 h before treatment with S1P (100 nM). Contraction was measured as indicated above. OGT inhibition was determined using the pan O-GlcNAc antibody anti-RL2.

**OGT knockdown with RNAi.**
Custom plasmids containing either mOGT siRNA (sequence: 5'-GAUUAAGCCUGUUGAAGUCTT-3') or scramble siRNA were purchased from Genescript. NIH3T3 cells were transfected using Lipofectamine RNAiMAX Transfection Reagent (ThermoFisher Scientific; 13778030) according to the manufacturer's protocol. After 24 h, cells were seeded at 1 x $10^5$ cells in 6-well dish for an additional 24 h before treatment with S1P (100 nM). Contraction was characterized as stated above. OGT knockdown was determined via Western blotting using anti-OGT to visualized OGT expression and anti-RL2 to visualized global O-GlcNAcylation.

**5SGlcNAc/DMSO/Thiamet-G S1P concentration course.**
Seeded NIH3T3 cells were treated with either DMSO vehicle, 5SGlcNAc (200 μM), or Thiamet-G (10 μM). After 20, 16, or 20 h respectively, wells were treated with S1P (0.05 - 5 μM) for 30 min. Contraction was characterized consistent with previously described experiments.

**Relaxation assay S1P time course.**
NIH3T3 cells were treated with either DMSO vehicle or 5SGlcNAc (200 μM) for 16 h before addition of S1P (100 nM). Images of each well were taken at 0, 30, 60, 120, and 180 min. Contraction was characterized as stated above.

**High/low glucose S1P concentration course.**
NIH3T3 cells were cultured in media containing either high (9 g $L^{-1}$) or low (1 g $L^{-1}$) concentrations of glucose for 24 h prior to being seeded as indicated above and incubated for an additional 24 h at which point cells were treated with S1P (0.05 - 5 μM) for 30 min. Contraction was characterized and changes in global O-GlcNAcylation were visualized using anti-RL2 as a pan O-GlcNAc antibody.

**S1PR1-4 antagonist screen.**
Seeded NIH3T3 cells were treated with 5SGlcNAc. After 16 h, cells were treated with one of the following; S1PR1 antagonist W146 (1 μM), S1PR2 antagonist JTE013 (1 μM), S1PR3 antagonist TY52156 (1 μM), S1PR4 antagonist CYM50358 (1 μM) for 10 min. Each well was then treated with S1P (100 nM) for 30 min and contraction was characterized as stated above.

**S1PR2 and S1PR5 agonist screen.**
Seeded NIH3T3 cells were treated with 5SGlcNAc. After 16 h, cells were treated with either S1PR2 agonist CYM5220 (1 or 5 μM) or S1PR5 agonist A971432 (1 μM) for 30 min. Contraction was characterized consistent with previously described experiments.

**S1PR2 knockdown with RNAi.**
NIH3T3 cells were seeded at 30% confluency in a 10 cm dish 24 h before transfection. 10 nmole of RNAi targeted against mouse S1PR2 or a scramble sequence was purchased from MilliporeSigma (siRNA ID: SASI_Mm01_00082880) and diluted in 1 mL in nuclease free $H_2O$ to make a stock concentration of 10 μM. Cells were transfected with 125 pmole (12.5 μL) of RNAi using Lipofectamine RNAiMAX transfection reagent (ThermoFisher Scientific) according to the manufacturer's protocol. 24 h post-transfection, cells were seeded 1 x $10^5$ cells in 6-well dishes. After 8 h, wells were treated with either DMSO vehicle or 5SGlcNAc (200 μM) for 16 h. Each well was then treated with S1P (100 nM) for 30 min and contraction was characterized as stated previously.

**ROCK1/2 inhibition with Y27632.**
NIH3T3 cells were seeded and treated with 5SGlcNAc (200 μM). After 15 h, cells were treated with ROCK1/2 inhibitor Y27632 (10 μM) for 1 h prior to treatment with S1P (0.05 - 5 μM) for 30 min. Contraction was characterized as stated above.

**MYPT1 vs MYPT1 mutants S1P concentration course.**
NIH3T3 cell-lines expressing human FLAG-tagged MYPT1, MYPT1(S/TtoA), MYPT1(d564-578), MYPT1(d588-602), or MYPT1Δ were transfected at 40% confluency with RNAi targeting mouse MYPT1 (transfection details described in subsequent section). After 24 h, cells were seeded 1 x $10^5$ cells in 6-well dishes and grown for an additional 24 h. After 48 h total, cell-lines were treated with S1P (0.05 or 0.1 μM) for 30 min. Contraction was characterized as stated above.

**MYPT1 vs MYPT1Δ S1P concentration course.**
NIH3T3 cell-lines expressing human FLAG-tagged MYPT1 or MYPT1Δ were transfected at 40% confluency with RNAi targeting mouse MYPT1 (transfection details described in a subsequent section). After 24 h, cells were seeded 1 x $10^5$ cells in 6-well dishes and grown for an additional 24 h. After 48 h total, cell-lines were treated with S1P (0.05 - 5 μM) for 30 min. Contraction was characterized as stated above.

**MYPT1 vs MYPT1Δ relaxation assay S1P time course.**
NIH3T3 cell-lines expressing human FLAG-tagged MYPT1 or MYPT1Δ were transfected at 40% confluency with RNAi targeting mouse MYPT1 (transfection details described in a subsequent section). After 24 h, cells were seeded 1 x $10^5$ cells in 6-well dishes and grown for an additional 24 h. After 48 h total, cell-lines were treated with S1P (100 nM). Each well was imaged at 0, 30, 60, and 120 min post S1P treatment. Contraction was characterized as stated above.

**MYPT1 vs MYPT1 + 5SGlcNAc vs MYPT1Δ S1P concentration course.**
Cell-lines expressing human FLAG-tagged MYPT1 or MYPT1Δ were transfected at 40% confluency with RNAi targeting mouse MYPT1 (transfection details described in a subsequent section). 24 h post transfection, cells were seeded 1 x $10^5$ cells in 6-well dishes. After 8 h, one plate of cells expressing FLAG-tagged MYPT1 was treated with 5SGlcNAc (200 μM) and one plate each of FLAG-tagged MYPT1 or MYPT1Δ were treated with DMSO vehicle. All three sets of cells were incubated for an additional 16 h before treatment with S1P (0.05 - 5 μM) for 30 min. Contraction was characterized as stated above.

**MYPT1Δ vs MYPT1Δ + Thiamet-G S1P concentration course.**
Cell-lines expressing human FLAG-tagged MYPT1Δ were transfected at 40% confluency with RNAi targeting mouse MYPT1 (transfection details described in a subsequent section). 24 h post transfection, cells were seeded 1 x $10^5$ cells in 6-well dishes for 8 h before treatment with either DMSO vehicle or Thiamet-G (10 μM). Cells were incubated for an additional 20 h at which point cells were treated with S1P (0.05 - 5 μM) for 30 min. Contraction was characterized as stated above.

**Human dermal fibroblast S1P concentration course.**
Human dermal fibroblast cell-line, BJ-5ta, were seeded 1.5 x $10^5$ cells in a 6-well dish. After 8 h, cells were treated with either DMSO vehicle or 5SGlcNAc (200 μM). Cells were incubated for an additional 16 h before treatment with S1P (0.05 - 5 μM) for 30 min. Contraction was characterized as stated above.

**Video characterization of S1P-mediated contraction.**
Seeded NIH3T3 cells were treated with either DMSO vehicle or 5SGlcNAc (200 μM) for 16 h. The following videos were taken using a Leica Microscope at 20x magnification: DMSO, DMSO + S1P (100 nM), DMSO + 10% FCS, 5SGlcNAc, 5SGlcNAc + S1P (100 nM), 5SGlcNAc + 10% FCS. For each null video, (DMSO or 5SGlcNAc), plates were placed under the microscope and recorded for 10 min. For wells treated with S1P or 10% FCS, video recording started immediately following treatment and continued for 10 min.

**Caspase-3 activation Western blot.**
NIH3T3 cells were treated with either DMSO vehicle or 5SGlcNAc (200 μM) for 16 h before the addition of S1P (100 nM) for 0, 30, 60, or 120 min at which point cells were collected by trypsinization and washed two times with PBS (2 min, 2,000 x g, 4 °C). Cell pellets were then resuspended in 4% SDS buffer (4% SDS, 150 mM NaCl, 50 mM TEA, pH 7.4) supplemented with 5 mg $mL^{-1}$ c0mplete mini protease inhibitor cocktail (MilliporeSigma), 5 mg $mL^{-1}$ PhosSTOP (MilliporeSigma), 1 mM phenylmethylsulfonyl fluoride. Cells were lysed via tip sonication (3x 5 sec on 5 sec off) and cell debris were pelleted (10 min, 10,000 x g, rt). Protein concentration was determined by BCA Assay (Pierce, ThermoScientific), and the lysate was diluted to 4 mg $mL^{-1}$ in lysis buffer before an equivalent volume of 2x loading buffer (20% glycerol, 0.2% bromophenol blue, 1.4% β-mercaptoethanol, pH 6.8). 20 μL (40 μg) per sample was loaded per lane. Western blotting was

performed according to the General Western blot procedure described above. Caspase-3 cleavage was visualized using anti-Caspase-3 and anti-Cleaved Caspase-3.

**RT-PCR confirming mRNA expression of S1PR1-5 in NIH3T3 cells.**
NIH3T3 cell lysate was harvested via trypsinization and washed twice with 1 mL cold PBS (5 min, 2,000 x g, 4 °C). RNA extraction was performed using RNeasy Minikit (Qiagen) according to manufacturer's protocol. RNA concentration was acquired using UV/Vis absorption and 1 µg RNA per reaction was subjected to RT-PCR. RT-PCR primers for each S1P receptor were designed with their mRNA sequences found using NCBI Gene Search tool (ncbi.nlm.nih.gov/gene). Sequences were copied into a RT-PCR primer design database (MIT Primer3). Primers were generated by accepting all design database default parameters and choosing primer pairs with the highest score. To verify sequence autonomy and avoid sequences that span mRNA introns, amplified region was copied into the UCSC genome browser and a BLAT search was performed. RT-PCR primers for S1PR1 (Forward: 5'-CTCCGGTTTCTTCTCACCCT-3', Reverse: 5'-TCCCAACAACTTGACCCAGT-3'), S1PR2 (Forward: 5'-GGTGCTGTCTGACCTCTTCT-3', Reverse: AAGGAGGACATCTGGGAAGC-3'), S1PR3 (Forward: 5'-ATCCAACCCTCACCCTGAAG-3', Reverse: AGGGAGAGAAGCAAGACCAC-3'), S1PR4 (Forward: 5'-GAGGGCAACTTGACGTGTTT-3', Reverse: 5'-ATTCAGAACAGGAAGGGCGA-3'), and S1PR5 (Forward: 5'-TGTGGGGAAGTCTTGTTGGT-3', Reverse: 5'-ACATCACCTGGTTCTGAGCA-3') were ordered along with primers for GAPDH mRNA (Forward: 5'-AGGCCGGTGCTGAGTATGTC-3', Reverse: TGCCTGCTTCACCACCTTCT-3') for use as loading controls (Integrated DNA Technologies). PCR reaction was carried out using Superscript IV One-Step RT-PCR System (Invitrogen) according to manufacturer's protocol. 10 µL from each PCR reaction was removed and added to fresh tubes containing 2 µL loading buffer (New England BioLabs). The entire 12 µL was loaded into a 1% agarose DNA gel (500 mg agarose, 5 µL ethidium bromide, 50 mL 1x Bio-Rad TAE buffer) and run for 45 min at 100 V. DNA bands were visualized using the ChemiDoc XRS+ molecular imager (Bio-Rad).

**MYPT1 and MLC phosphorylation Western blot time course.**

***General procedure.*** NIH3T3 or stable cell-lines were seeded 5 x $10^5$ cells per 10 cm dish. To each plate, S1P at indicated concentration was added and plates were incubated for 0, 2, 5, 10, or 30 min. At each time point, dishes were removed from the incubator, media was decanted, and plates were immediately placed on ice, rinsed once with ice cold PBS, and harvested by scraping into pre-cooled 15 mL falcon tubes. Cells were pelleted by centrifugation (5 min, 2,000 x g, 4 °C). Pellets were resuspended in 4% SDS buffer (4% SDS, 150 mM NaCl, 50 mM TEA, pH 7.4) and lysed via tip sonication (3x 5 sec on 5 sec off). Lysate was centrifuged (10 min, 10,000 x g, rt) and supernatant was moved to a fresh tube. Protein concentration was determined by BCA Assay (Pierce, ThermoScientific) and gel samples were prepared at 2 mg $mL^{-1}$ with appropriate volumes of 4% SDS buffer and 2x loading buffer (20% glycerol, 0.2% bromophenol blue, 1.4% β-mercaptoethanol, pH 6.8). 20 µL (40 µg) per sample was loaded per lane. Western blotting was performed according to the General Western blot procedure described above. Induction of MYPT1 phosphorylation was visualized using anti-pT696 MYPT1 and anti-p853 MYPT1. Activation of MLC was visualized using anti-pT18/S19 MLC2. Expression of MYPT1 and MLC2 were used as loading controls.

***MLC activation with high S1P signaling.*** To validate S1P's role in mediating MLC activation, NIH3T3 cells were treated with a high concentration S1P (5 µM) for 0, 2, 5, 10, and 30 min, and worked up as indicated in the previous section. Activation of the pathway was visualized with Western blotting against pT18/S19 MLC and MLC2.

***O-GlcNAcylation sensitizes S1P-mediated phosphorylation of MLC and MYPT1.*** To indicate the role O-GlcNAcylation plays in MLC activation, 8 h after plating, cells were treated with DMSO vehicle or 5SGlcNAc (200 µM) and incubated for 16 h. To each plate per original treatment (DMSO or 5SGlcNAc) low S1P (100 nM) was added and plates were incubated for 0, 2, 5, 10, and 30 min and worked up as described previously. Induction of MYPT1 phosphorylation was visualized using anti-pT696 MYPT1 and anti-p853 MYPT1. Activation of MLC was visualized using anti-pT18/S19 MLC2. Expression of MYPT1 and MLC2 were used as loading controls.

***MYPT1 vs MYPT1Δ phosphorylation Western blot time course.*** Stable expressing human Wild-Type or Δ550-600 MYPT1 constructs were seeded at 30% confluency 24 h before transfection with RNAi targeting mouse MYPT1. After 24 h, cells were seeded 5 x $10^5$ cells per 10 cm dish and grown for an additional 24 h at which point they were treated with a low concentration of S1P (100 nM) for 0, 2, 5, or 10 min. Cells were worked up as described above. Induction of MYPT1 phosphorylation was visualized using anti-pT696 MYPT1 and anti-p853 MYPT1. Activation of MLC was visualized using anti-pT18/S19 MLC2. Expression of MYPT1 and MLC2 were used as loading controls.

***MYPT1Δ MLC activation time course.*** To ensure that MYPT1Δ continues to perform its biological functions, NIH3T3 cells expressing FLAG-tagged MYPT1Δ were seeded at 30% confluency 24 h before transfection with RNAi targeting mouse MYPT1. After 24 h, cells were seeded 5 x $10^5$ cells per 10 cm dish and grown for an additional 24 h at which point they were treated with S1P (100 nM) for 0, 60, and 180 min. Cells were worked up as described above. Activation of MLC was visualized using anti-pT18/S19 MLC2. Expression MLC2 was used as a loading controls.

**Expression and purification of GalT(Y289L).**

pET23a GalT(Y289L) plasmid was provided by P. Qasba, National Cancer Institute. The plasmid was transformed in BL21 *E. Coli* (Novagen). A 1 L culture containing ampicillin (100 μg $mL^{-1}$) was inoculated with 10 mL from a 50 mL starter culture grown overnight at 37 °C. 1 L cultures were grown at 37 °C until an OD (A600) of 0.60 was obtained. At this point, expression was induced using isopropyl β-D-1-thiogalactopyranoside (1 mM final concentration, 1000x stock in $H_2O$) for 4 h at 37 °C. Bacterial cells were harvested by centrifugation (10 min, 4,000 x g, 4 °C) and resuspended in 10 mL of suspension buffer (25% sucrose w/v in 1x PBS). Cells were lysed via tip sonication (30 sec on 30 sec off, 12 min total, 4 °C). The resulting lysate was diluted to 80 mL in cold suspension buffer. Inclusion bodies were harvested by centrifugation (30 min, 15,000 x g, 4 °C) and washed in cold suspension buffer followed by centrifugation until suspension turned white (approximately 8 - 10 times). Inclusion bodies were then washed once with cold wash buffer (10 mM phosphate, pH 7.0) and harvested via centrifugation. Inclusion bodies were resuspended in 14 mL of cold $H_2O$ and poured over solid guanidine HCl and $Na_2SO_3$ resulting in a solution of 5 M GuHCl and 300 mM $Na_2SO_3$. The suspension was vortexed vigorously and the volume was adjusted to 25 mL with cold $H_2O$. Freshly made NTSB solution (50 mM DTNB, 1 M $Na_2SO_3$ in $H_2O$, pH 8.0) was added followed by vigorous vortexing in order to sulfenate free thiols. Completion of the reaction is indicated by a color change from dark orange to pale yellow. Protein was precipitated with 250 mL of cold $H_2O$ and centrifuged immediately (30 min, 9,900 x g, 4 °C). Protein pellets were washed 3x by resuspension in cold $H_2O$. Pellets were then resuspended in 14 mL cold $H_2O$ and poured over solid guanidine HCl, vigorously vortexed, and diluted to 25 mL in cold $H_2O$ (5 M GuHCl). Protein suspension was then diluted to a final concentration of 1 mg $mL^{-1}$ with additional 5 M GuHCl solution (OD (A275) ≈ 2.0). This solution was diluted 10-fold into cold refolding buffer (5 mM EDTA, 4 mM cysteamine, 2 mM cystamine, 100 mM Tris, pH 8.0) with gentle stirring and allowed to refold (48 h, 4 °C) without agitation. Refolded protein was dialyzed two times into fresh cold $H_2O$ (24 h, 4 °C) and concentrated using centrifugal filters (30 kDa cutoff, Amicon Ultra, MilliporeSigma) to 3 mL. Buffer was exchanged with 10 mL of reaction buffer (10 mM Tris, pH 8.0) and a final protein concentration was determined (OD (A275) ≈ 1.5). Purified protein was stored at 4 °C.

**General procedure for chemoenzymatic labeling.**

NIH3T3 cells were collected by trypsinization and washed two times with PBS (2 min, 2,000 x g, 4 °C). Cell pellets were then resuspended in 4% SDS buffer (4% SDS, 150 mM NaCl, 50 mM TEA, pH 7.4) supplemented with 5 mg $mL^{-1}$ c0mplete mini protease inhibitor cocktail (MilliporeSigma), 5 mg $mL^{-1}$ PhosSTOP (MilliporeSigma), 1 mM phenylmethylsulfonyl fluoride. Cells were lysed via tip sonication (3x 5 sec on 5 sec off) and cell debris were pelleted (10 min, 10,000 x g, rt). Protein concentration was determined by BCA Assay (Pierce, ThermoScientific), and the lysate was diluted to 1 mg $mL^{-1}$ in 1% SDS chemoenzymatic buffer (1% SDS, 20 mM HEPES, pH 7.9). Proteins were precipitated by adding a 3x volume of MeOH, a 0.75x volume of $CHCl_3$, and 2x volume of $H_2O$ followed by vortexing and centrifugation (5 min, 13,000 x g, rt). The aqueous phase was discarded without disturbing the interface layer before adding 2.5x volume of MeOH, vortexing briefly, and centrifugation (5 min, 13,000 x g, rt). The resulting protein pellet was allowed to air-dry for 5-10 min before being resuspended in 1% SDS chemoenzymatic buffer. Protein concentration was normalized using the BCA Assay (Pierce, ThermoScientific) and diluted to 2.5 mg $mL^{-1}$ in 1% SDS chemoenzymatic buffer. To start the chemoenzymatic transfer reaction, the following reagents were added in order for 100 μg of total protein: 49 μL of $H_2O$, 80 μL of labeling buffer (2.5x; 5% NP-40, 125 mM NaCl, 50 mM HEPES, pH 7.9), 55 μL of $MnCl_2$ (100 mM in $H_2O$), and 50 μL of UDP-GalNAz (0.5 mM in 10 mM HEPES, pH 7.9). This was mixed by pipetting gently. Finally, 7.5 μL of purified GalT(Y289L) (in 10 mM Tris, pH 8.0). The reaction mixture was incubated for 16 h at 4 °C without agitation. Following incubation, unreacted UDP-GalNAz was removed by MeOH/$CHCl_3$/$H_2O$ precipitation. Air-dried protein pellets were resuspended in 1% SDS CuAAC buffer (1%

SDS, 150 mM NaCl, 50 mM TEA, pH 7.4) and subjected to either immunoprecipitation or conjugation with DBCO-PEG mass-tags.

**Endogenous MYPT1 and MLC chemoenzymatic/biotin IP.**
NIH3T3 cells at 80% confluency were harvested, resuspended in 4% SDS buffer (4% SDS, 150 mM NaCl, 50 mM TEA, pH 7.4) supplemented with 5 mg $mL^{-1}$ c0mplete mini protease inhibitor cocktail (MilliporeSigma), 5 mg $mL^{-1}$ PhosSTOP (MilliporeSigma), 1 mM phenylmethylsulfonyl fluoride, and lysed via tip sonication (3x 5 sec on 5 sec off) before being subjected to chemoenzymatic transfer according to the protocol described above. Following chemoenzymatic transfer, proteins were precipitated by adding a 3x volume of MeOH, a 0.75x volume of $CHCl_3$, and 2x volume of $H_2O$, and resuspended in 1% SDS buffer (1% SDS, 150 mM NaCl, 50 mM TEA, pH 7.4). Inputs were generated by adding 25 µL of the lysis buffer and 25 µL of 4x LB (200 mM Tris, 8% SDS, 40% glycerol, 0.4% bromophenol blue, 2.8% β- mercaptoethanol, pH 6.8) to 50 µL of lysate.

**CuAAC enrichment.**
Cell lysates (1 mg at 1 mg $mL^{-1}$) labeled chemoenzymatically were subjected to CuAAC enrichment performed using a freshly made master mix containing alkyne-azo-biotin (100 µM from 5 mM stock in DMSO, Click Chemistry Tools), TCEP (1 mM from 50 mM freshly prepared stock in $H_2O$), TBTA (100 µM from 10 mM stock in DMSO), $CuSO_4{:}5H_2O$ (1 mM from 50 mM freshly prepared stock in $H_2O$). After 1 h incubation in the dark, proteins were precipitated by addition of 4x volume of ice-cold MeOH and incubation at -20 °C for 2 h. Proteins were collected by centrifugation (30 min, 5,000 x g, 4 °C) and washed 3x with ice-cold MeOH. Pellets were then air-dried for 15 min before being resuspended in 800 µL resuspension buffer (6 M urea, 2 M thiourea, 10 mM HEPES, pH 8.0). To ensure complete resuspension, mixture was incubated in a bath sonicator for 5 min. Streptavidin beads (25 µL of a 50% slurry per sample, ThermoFisher Scientific) were allocated into 2 mL dolphin-nosed tubes and prepared by washing 2x with 1 mL PBS and 1x with 1 mL resuspension buffer before a final resuspension in 200 µL resuspension buffer. Samples were added to the dolphin-nosed tubes and incubated on a rotator (2 h, rt, full rotation). Beads were washed 2x with 1 mL resuspension buffer, 2x with 1 mL PBS, and 2x with 1 mL 1% SDS in PBS. Beads were then incubated in 25 µL of sodium dithionite solution (1% SDS, 25 mM sodium dithionite) for 30 min at rt to elute bound proteins. Beads were centrifuged (2 min, 2,000 x g, rt) and the eluent was collected. Elution was repeated and the two elution products were combined. Proteins were precipitated in 1 mL ice-cold MeOH overnight at -20 °C. Proteins were collected by centrifugation (10 min, 10,000 x g, 4 °C). Pellets air-dried for 5 min before a final resuspension in 30 µL of 4% SDS buffer (4% SDS, 150 mM NaCl, 50 mM TEA, pH 7.4), and bath sonicated to ensure complete dissolution. Gel samples were prepared by adding 30 µL of 2× loading buffer (20% glycerol, 0.2% bromophenol blue, 1.4% β-mercaptoethanol) and boiling (5 min, 98 °C). Inputs and IP samples were separated by SDS-PAGE and Western blotted for Nup62 (positive control), β-actin (negative control), MYPT1, and MLC2 according to the procedure described above.

**Mass-shift of MYPT1.**
NIH3T3 cells were plated in 2x 150 mm dishes and grown to 80% confluency. Cells were harvested by trypsinization and washed two times PBS (2 min, 2,000 x g, 4 °C). The resulting cell pellets were resuspended in 500 µL 4% SDS lysis buffer (4% SDS, 10 mM TEA pH 7.4, 150 mM NaCl) supplemented with 5 mg $mL^{-1}$ c0mplete mini protease inhibitor cocktail (MilliporeSigma), 5 mg $mL^{-1}$ PhosSTOP (MilliporeSigma), 1 mM phenylmethylsulfonyl fluoride. The resulting suspension was lysed using a tip sonicator (3x 5 sec on 5 sec off). 5 µL of TCEP (500 mM stock dissolved in 1 M NaOH) was then added to each sample and boiled at (10 min, 98 °C). Samples were cooled to room temperature before adding 40 µL iodoacetimade (600 mM stock dissolved in 4% SDS buffer) and incubated in the dark for 30 min. Samples ere diluted to a total volume of 2 mL using 1% SDS buffer (1% SDS, 50 mM TEA pH 7.4, 150 mM NaCl). Proteins were precipitated by adding 3x volume of MeOH, 0.75x volume of $CHCl_3$, and 2x volume of $H_2O$ and briefly vortexing followed by centrifugation (5 min, 5,000 x g, rt). The aqueous phase was discarded without disturbing the interface between layers before adding a 2.5x volume of MeOH, vortexing, and pelleting the protein (10 min, 5,000 x g, rt). Pellets were washed with MeOH (3x 1 mL). The resulting pellet was allowed to air-dry for 5 min before being resuspended in 400 µL 1% SDS chemoenzymatic buffer (1% SDS, 20 mM HEPES, pH 7.9). Protein concentration was normalized using the BCA Assay and diluted to 2.5 mg $mL^{-1}$ in 1% SDS chemoenzymatic buffer. UDP-GalNAz Enzymatic Labeling was set up and scaled as described above. For each sample, 200 µg of protein lysate was incubated with both GalT and UDP-GalNAz. Negative controls for each sample, incubating with just GalT, were generated using 100 µg of protein lysate. After chemoenzymatic labeling (protocol described above), unreacted UDP-GalNAz was removed via $MeOH/CHCl_3/H_2O$ precipitation. Air-dried protein pellets were resuspended in 90 µL (negative samples) or 180 µL (positive samples) of 1% SDS

(1% SDS, 150 mM NaCl, 50 mM TEA, pH 7.4). 10 µL (negative samples) or 20 µL (positive samples) of 10 mM DBCO-PEG5000 or DBCO-PEG2000 in DMSO were added and the mixture was boiled (5 min, 98 °C). Unreacted DBCO-PEG was removed by MeOH/$CHCl_3$/$H_2O$ precipitation described previously. The pellet was again to air-dried for 5-10 min. Finally, 25 µL 4% SDS buffer was added to each sample. The mixture was briefly sonicated in a bath sonicator to ensure complete resuspension before 25 µL of 2x loading buffer (20% glycerol, 0.2% bromophenol blue, 1.4% β-mercaptoethanol, pH 6.8) was added and the samples were boiled (5 min, 98 °C). 20 µL (40 µg) of each gel sample was loaded per lane. Western blots were performed according to the General Western blot procedure described above with the following two changes: anti-MYPT1 primary antibody was incubated at a dilution of 1:500 and HRP-conjugated secondary antibodies were incubated at a dilution of 1:5,000.

**FLAG-IP for RL2 detection of MYPT1.**

FLAG-tagged MYPT1 expressing NIH3T3 grown to 80% confluency were harvested by trypsinization and washed two times with PBS (2 min, 2,000 x g, 4 °C) Cell pellets were then resuspended in 200 µL of lysis buffer (1% Triton-x 100, 50 mM Tris, 150 mM NaCl, 1 mM EDTA) supplemented with 5 mg $mL^{-1}$ c0mplete mini protease inhibitor cocktail (MilliporeSigma), 5 mg $mL^{-1}$ PhosSTOP (MilliporeSigma), 1 mM phenylmethylsulfonyl fluoride, and 100 µM Thiamet-G. This suspension was lysed using a tip sonicator (3x 5 sec on 5 sec off) on ice. The suspension was then centrifuged (5 min, 15,000 x g, 4 °C) to isolate the soluble lysate, disposing of the remaining pellet. Protein concentration was determined by BCA Assay and diluted to 2 mg $mL^{-1}$ in lysis buffer. Inputs were generated by adding 25 µL of the lysis buffer and 25 µL of 4x LB (200 mM Tris, 8% SDS, 40% glycerol, 0.4% bromophenol blue, 2.8% β- mercaptoethanol, pH 6.8) to 50 µL of lysate. 20 µL of Anti-FLAG-M2 magnetic beads (MilliporeSigma) were added to an eppendorf tube and washed with lysis buffer (3x 1 mL). Lysate was diluted to 1 mg $mL^{-1}$ in lysis buffer. 1 mL of lysate was added to the beads followed by incubation for 1 h rotating at 4 °C. Beads were collected and washed (6x 1 mL lysis buffer). To elute from beads, 30 µL of lysis buffer and 10 µL of 4x LB were added to beads, vortexed briefly, and boiled (5 min, 98 °C). Beads were pelleted by centrifugation (1 min, 15,000 x g, rt). The resulting solution was transferred to a new eppendorf tube. For detection was anti-FLAG, 5 µL of input and 10 µL of IP was loaded. For detection of RL2 15 µL of input and 30 µL of IP was loaded. Western blot was performed according to General Western blot Procedure described above.

**FLAG-IP for RL2 detection of MYPT1 vs MYPT1Δ.**

FLAG-tagged MYPT1 or MYPT1Δ NIH3T3 cell-lines were harvested and worked-up as described in the previous section. Western blot was performed according to General Western blot Procedure described above.

**Co-Immunoprecipitation of FLAG-tagged MYPT1 or FLAG-tagged MYPT1Δ and ROCK1/2.**

NIH3T3 cells were treated with DMSO vehicle or 5SGlcNAc (200 µM) for 16 h before being harvested by trypsinization. Cell lysate was washed twice with 1 mL cold PBS (5 min, 2,000 x g, 4 °C) and resuspended in 500 µL lysis buffer (1% IGPAL, 50 mM TEA, 150 mM NaCl, pH 7.4) supplemented with 5 mg mL-1 c0mplete mini protease inhibitor cocktail (MilliporeSigma), 5 mg mL-1 PhosSTOP (MilliporeSigma), 1 mM phenylmethylsulfonyl fluoride. The cell suspension was lysed using a tip sonicator (3x 5 sec on 5 sec off) and any remaining cell debris were pelleted (10 min, 10,000 x g, 4 °C). Lysate was moved to a pre-chilled tube and kept on ice. Protein concentration was determined by BCA Assay (Peirce, ThermoScientific) and diluted to 2 mg mL-1 in lysis buffer. Inputs were generated by adding 25 µL of the lysis buffer and 25 µL of 4x LB (200 mM Tris, 8% SDS, 40% glycerol, 0.4% bromophenol blue, 2.8% β- mercaptoethanol, pH 6.8) to 50 µL of lysate. An anti-flag co-immunoprecipitation was performed using the Catch and Release system (ThermoScientific) according to the manufacturers protocol. Briefly, spin columns were washed twice with 400 µL 1x Wash Buffer (30 sec, 2000 x g, 4 °C). 500 µg of cell lysate (250 µL) per sample was added to designated spin columns followed by 5 µL PMSF (200 µM stock in IPA), 225 µL 1x Wash Buffer, 10 µL Affinity Ligand, and 10 µL of anti-FLAG antibody. Plugged spin columns were then incubated on a rotator (1 h, 4 °C, full rotation). Spin columns were next centrifuged (30 sec, 2000 x g, 4 °C) and washed 3x with 1x Wash Buffer (30 sec, 2000 x g, 4 °C) to remove unbound proteins. Spin columns were then placed in fresh capture tubes and bound proteins were eluted in 70 µL of 1x Denaturing Elution buffer containing βME to generating gel samples. Input and IP gel samples were boiled (5 min, 95 °C) and separated by SDS-PAGE. ROCK enrichment was detected by Western blotting and normalized to overall protein capture via Coomassie staining. The results were quantified and presented as mean ± SEM of the normalized ROCK levels (n = 3 biological replicates). Statistical significance was determined using a 2-tailed, unpaired Student's t-test.

**Generation of NIH3T3 cell-lines stably expressing FLAG-tagged MYPT1 mutants.**
FLAG-tagged MYPT1, MYPT1(S/TtoA), MYPT1(d564-578), MYPT1(d588-602), and MYPT1Δ NIH3T3 cell-lines were generated using the PiggyBac™ Transposon Vector System (System Biosciences). Briefly, NIH3T3 cells at 30% confluency in 10 cm dishes were co-transfected with 5 μg PiggyBac transposase promoter (plasmid ID: PB531A-1) and 10 μg PiggyBac transposon plasmid pPB-CAG-IRES2-puro containing either human MYPT1, human MYPT1(S/TtoA), human MYPT1(d564-578), human MYPT1(d588-602), or human MYPT1Δ550-600 (MYPT1Δ). One plate was transfected with 10 μg of pcDNA3 to serve negative selection control. 24 h post-transfection, plates were split 1:2 into normal growth media (DMEM + 10% FCS) and supplemented with 1 μg $mL^{-1}$ puromycin (1 mg $mL^{-1}$ stock in $H_2O$). Cells were selected for three days at which point puromycin concentration was reduced to 250 ng $mL^{-1}$. FLAG-tagged MYPT1 expression was confirmed with analysis by Western blotting against anti-FLAG and anti-MYPT1. Effect of MYPT1 vs MYPT1Δ on global O-GlcNAcylation and OGT expression were analyzed by Western blotting against anti-RL2 and anti-OGT.

**Collagen gel contraction assay.**
3D collagen contraction assay was performed using a two-step cell contraction assay kit purchased from Cell Biolabs (CBA-201) according to manufacturers protocol. Briefly, BJ-5ta cells was seeded 2 x $10^5$ used per well. A 4:1 mixture of collagen stock solution to cells was mixed in a 15 mL falcon tube and 500 μL per well was plated in a 24-well dish. The cell-collagen matrix was incubated at 37 °C for 1 h to allow collagen polymerization before adding 1 mL of media to each well. After 30 hours, wells were supplemented with either DMSO vehicle or 5SGlcNAc inhibitor (200 μM) and incubated for another 16 hours at which point the media was changed to serum-free media containing S1P (0.1 - 5 μM). Contraction was initiated by gently releasing the sides and bottom of the matrices from the plate. Images were taken immediately after media change (t=0) and after 30 min using the ChemiDoc XRS+ molecular imager (Bio-Rad). Contraction was measured using ImageJ. Specifically, collagen matrix diameter was measured by taking the average of one horizontal (0 degrees) and one vertical (90 degrees) measurement of each matrix.

**Endogenous mouse MYPT1 knockdown with RNAi.**
NIH3T3 or stably expressing cell-lines were seeded at 30% confluency in 10 cm dishes 24 h before transfection. 5 nmole of RNAi targeted against mouse MYPT1 or a scramble sequence was purchased from ThermoFisher Scientific (siRNA ID: s70342) and diluted in 500 μL in nuclease free $H_2O$ to make a stock concentration of 10 μM. Cells were transfected with 125 pmole (12.5 μL) of RNAi using Lipofectamine RNAiMAX transfection reagent (ThermoFisher Scientific) according to the manufacturer's protocol. Knockdown efficiency was measured via Western blotting against MYPT1. Western blot procedure is described in detail previously.
